# Necessity of integrated genomic analysis to establish a designed knock-in mouse from CRISPR-Cas9-induced mutants

**DOI:** 10.1101/2022.06.08.495409

**Authors:** Masahide Yoshida, Tomoko Saito, Yuki Takayanagi, Yoshikazu Totsuka, Tatsushi Onaka

## Abstract

The CRISPR-Cas9 method for generation of knock-in mutations in a rodent embryo yields many F0 generation candidates that may have the designed mutations. Genetic mutations must be confirmed using genomic DNA with a mixture of designed and unexpected mutations. In our study for establishing Prlhr-Venus knock-in reporter mice, we found that genomic rearrangements near the targeted knock-in allele, tandem multicopy at a target allele locus and mosaic genotype of two different knock-in alleles occurred in addition to the designed knock-in mutation in the F0 generation. Conventional PCR and genomic sequencing were not able to detect mosaicism or discriminate between the designed one-copy knock-in mutant and a multicopy-inserted mutant. However, by using a combination of Southern blotting and the next-generation sequencing-based RAISING method, these mutants were successfully detected in the F0 generation. In F1 and F2 generations, droplet digital PCR analysis contributed to establishment of the strain, although a multicopy was falsely detected as one copy by the analysis in the F0 generation. We emphasize that the combination of these five methods allowed us to select promising F0 generations and establish the designed strain and that focusing only on positive evidence of knock-in can lead to erroneous selection of undesirable strains.

## Introduction

Gene targeting, which transduces a mutation in a specific endogenous gene, has been broadly used to generate animal models for clarification of physiological or pathological functions. Homologous recombination in embryonic stem (ES) cells has been classically used to obtain gene-targeted rodents (Capecchi, 2005). However, by using this method, it takes nearly one year to obtain a genetically targeted mouse including the time for producing correctly targeted ES cell clones and acquiring animals capable of reproduction from chimeric mice. On the other hand, genome editing methods with zinc finger nuclease (ZFN) (Geurts et al., 2009), transcription activator-like effector nucleases (TALEN) (Miller et al., 2011) and CRISPR-Cas9 ribonucleoprotein complexes (Cong et al., 2013; Mali et al., 2013) are promising approaches for obtaining gene-targeted rodents in a limited period of time and for accelerating biological and medical research. The CRISPR-Cas9 method has been developed remarkably because of its simplicity and efficiency (Wang et al., 2013). In homologous recombination in ES cells, a gene targeting vector is electroporated into ES cells and drug-resistant ES cells are screened. Southern blotting or PCR analysis is used to further select ES cell clones that have been recombined correctly. The ES cell clones are microinjected into blastocysts to obtain germline-transmitted chimeric rodents (Robertson, 1991). On the other hand, in the CRISPR-Cas9 method, the combination of CRISPR RNA (crRNA) including the recognition sequence complementary to the targeting genomic sequence and trans-activating CRISPR RNA (tracrRNA), or single guide RNA (sgRNA), and Cas9 nuclease are co-injected into fertilized eggs and the target gene is directly modified within embryos (Doudna and Charpentier, 2014; Shen et al., 2013). While it takes only three months to obtain adult candidate rodents, the genetic variation must be investigated using genomic DNA extracted from somatic tissues of individual rodents.

In the past few years, targeted knock-in methods have been developed by administering a donor DNA template along with a component of the CRISPR-Cas9 complex. The CRISPR-Cas9 complex recognizes a specific sequence and cleaves double-stranded DNA (dsDNA) adjusted to the PAM sequence. In mammalian cells, site-specific dsDNA breaks are repaired by nonhomologous end joining (NHEJ), homology-directed repair (HDR) or microhomology-mediated end joining (MMEJ) mechanisms (Patterson-Fortin and D’Andrea, 2020; Wang and Xu, 2017). Knock-in events can occur when repair of dsDNA breaks is accomplished using a donor DNA template with two homology arms on either side of the transgene of interest (Xue and Greene, 2021). Many researchers in the genome editing community have been endeavoring to improve the efficiency and specificity of CRISPR-Cas9-based knock-in methods in rodent embryos (Gu et al., 2018; Lin et al., 2018; Ma et al., 2014; Quadros et al., 2017; Yang et al., 2013; Yao et al., 2017; Yao et al., 2018; Yoshimi et al., 2016; Yoshimi et al., 2021). However, several obstacles remain in producing an accurate and rapid knock-in rodent model using these techniques.

It has recently been reported that tandem multiple integration at a target locus is frequently observed in F0 generation mice that had been injected with the combination of two guide RNAs and one donor DNA containing two loxP sequences into the zygote for generation of conditional knock-out mice. In addition, the mosaic genotype of the F0 generation has been revealed by analysis of F1 generation mice (Codner et al., 2018; Skryabin et al., 2020). It was also pointed out that conventional PCR analysis, which is frequently used for genotype confirmation, failed to identify such multiple integration events in most cases, leading to a high rate of erroneous identification as correct recombination with a single copy (Skryabin et al., 2020).

The knock-in method using CRISPR-Cas9 can contribute to the insertion of reporter genes and the generation of preclinical rodent models such as models of triplet disease and cancer by insertion of causative sequences (Budworth and McMurray, 2013; Gurumurthy and Lloyd, 2019; Lampreht Tratar et al., 2018). It is fundamental and essential to obtain the designed and precise knock-in rodents in order to reproduce human pathological conditions and to elucidate the molecular mechanisms underlying diseases and physiological functions. In the CRISPR-Cas9 method, many candidate rodents that may have the designed mutation can be obtained (Quadros et al., 2017; Yao et al., 2018). However, analysis of the F1 generation by mating all candidate F0 generations results in unnecessary breeding of research rodents and requires a lot of time, effort, and space, which will be a bottleneck for the CRISPR-Cas9 method. Whole genome sequencing seems to be the best method to validate the results of genome editing, but it has been pointed out that whole genome sequencing is not effective in mosaic animals (Bunton-Stasyshyn et al., 2021). Although there are several methods for analyzing the genome structure, the only way to find rodents with the required knock-in allele from a mixture of designed and unpredictable mutants is to combine multiple methods. However, it is not clear which combination of methods is effective for identifying possible mutations produced by the CRISPR-Cas9-based knock-in method and for establishing a strain line that has the designed mutation.

In this study, we generated prolactin-releasing peptide receptor (Prlhr)-Venus knock-in reporter mice by a CRISPR-Cas9 method with one guide RNA and one long single-stranded DNA (lssDNA) making use of HDR mechanisms. We conducted five genome structural analyses including the recently developed Rapid Amplification of Integration Sites without Interference by Genomic DNA contamination (RAISING) method and compared the results over three generations to establish a strain. These analyses revealed that genomic rearrangement in the vicinity of the targeted knock-in allele, multicopy and mosaic genotype occurred in the F0 generation, and the complex genotypes were able to be detected by a combination of multiple methods.

## Results

### Generation of CRISPR-Cas9-based Prlhr-Venus knock-in mice with lssDNA and one sgRNA

A plan was made to generate Prlhr-Venus knock-in mice by inserting a Venus-SV40 polyadenylation signal cassette into the gene encoding Prlhr (Fig. 1A). The target site of sgRNA was 24 bp downstream of the Prlhr initiation codon to produce 9 amid acids of Prlhr and Venus fusion protein (Fig. 1B). We microinjected a mixture of human codon-optimized Cas9 nuclease (hCas9) mRNA, sgRNA and lssDNA into 347 embryos and transferred 334 two-cell embryos into pseudopregnant female mice. As a result, we obtained 42 pups of the F0 generation from 11 mothers (Fig. 1C).

**Fig. 1.**
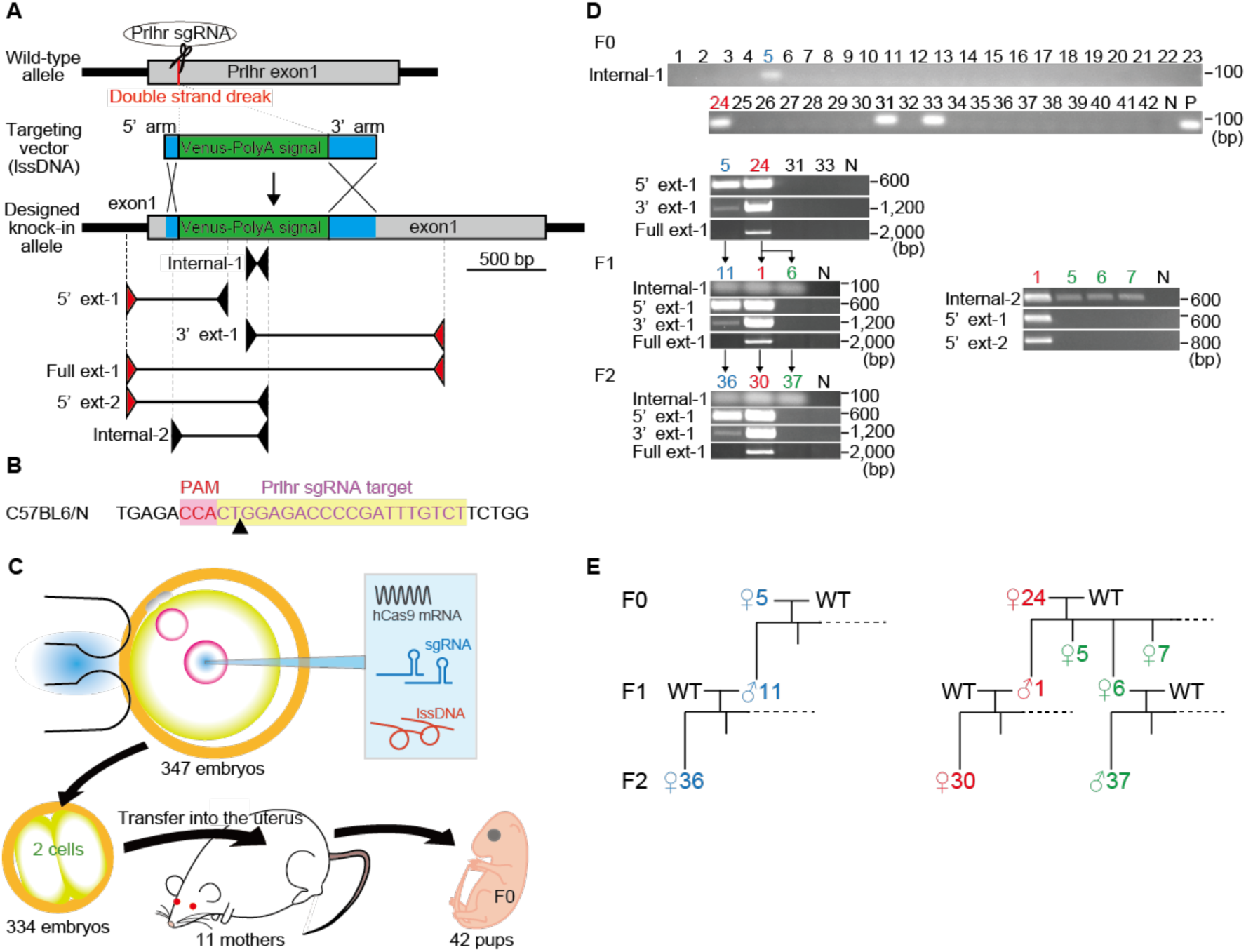
CRISPR-Cas9-mediated knock-in strategy at the prolactin releasing-peptide receptor (Prlhr) locus and analysis of Venus integration by using conventional PCR. (A) Schematic representation of wild-type mouse Prlhr genomic locus, sgRNA targeting site, targeting vector containing the Venus - polyadenylation signal and primer sets (internal primer pair, Internal-1and Internal-2: primer external to the targeting vector and internal primer pairs, 5’ext-1, 3’ext-1, 5’ext-2: primer pair external to the targeting vector, Full ext-1). Red and black arrows show the primer external to the targeting vector and internal primer, respectively. (B) Target sequence of Prlhr sgRNA on the C57BL6/N genome. (C) Schematic representation of hCas9 mRNA, sgRNA and lssDNA co-injection into mouse embryos to generate founders (F0) with the knock-in mutation. (D) Conventional PCR analysis of genomic DNA from F0, F1 and F2 generations by using each primer pair. Black arrows show the parent-offspring relationships. (E) Family trees of numbers 5 and 24 of the F0 generation.

### Conventional PCR analysis

When using the CRISPR-cas9 method for gene targeting in embryos, conventional PCR for the F0 generation is the initial step because it is convenient and sensitive. We first performed PCR analysis with an internal primer pair within the Venus-SV40 polyadenylation signal cassette. The PCR analysis revealed that 4 of the 42 pups obtained showed positive amplification (upper panel of Fig. 1D, Internal-1: numbers 5, 24, 31 and 33 of F0). We then used primers external to the targeting vector and internal primers in combination. PCR products of the 5’ and 3’ regions representing a potentially targeted allele were detected in numbers 5 and 24 of F0 (second panel of Fig. 1D, 5’ ext-1 and 3’ ext-1). The PCR product by use of the primer pair external to the targeting vector was only found in number 24 of F0 (second panel of Fig. 1D, Full ext-1).

We crossed F0 numbers 5 and 24 with the wild type to obtain the F1 generation. Number 11 of F1 was the offspring of number 5 of F0 and numbers 1, 5, 6 and 7 of F1 were offsprings of number 24 of F0 (Fig. 1E). The PCR product of the internal primer pair was detected in numbers 11, 1 and 6 of F1 (left third panel of Fig. 1D, Internal-1). PCR products of the 5’ and 3’ regions representing a potentially targeted allele were detected in numbers 11 and 1 but not in number 6 of F1 (left third panel of Fig. 1D, 5’ ext-1 and 3’ ext-1). The PCR product by use of the primer pair external to the targeting vector was found in number 1 of F1 (left third panel of Fig. 1D, Full ext-1). We additionally analyzed offsprings of number 24 of F0 and numbers 1, 5, 6 and 7 of F1 by using other primer pairs. We found PCR products in all 4 offsprings by using the internal primer pair (numbers 1, 5, 6 and 7 of F1). However, PCR products of the 5’ regions by using the external-internal primer pairs were not detected in numbers 5, 6 and 7 of F1 (right third panel of Fig. 1D, Internal-2, 5’ext -1 and 2). We crossed F1 numbers 11, 1 and 6 with the wild type to obtain the F2 generation. Number 36 of F2 was the offspring of number 11 of F1. Number 30 of F2 was the offspring of number 1 of F1 and number 37 was offspring of number 6 of F1 (Fig. 1E). The detection pattern of PCR products in F2 was exactly same as that of their parents (fourth panel of Fig. 1D).

Two candidate F0 generation mice with Venus insertion in the Prlhr locus were detected by conventional PCR. On the other hand, analysis of their F1 and F2 generation progenies showed that F0 number 24 was a mosaic with at least two kinds of Venus insertions that can be separated by PCR analysis.

### Droplet digital PCR analysis

We conducted droplet digital PCR to determine the copy number of the transgene. We first confirmed whether the copy number of the Venus gene was accurately detected by using droplet digital PCR. Oxytocin receptor-Venus knock-in heterozygous mice generated by using ES cells were used for a positive control as a single copy of the Venus gene (Yoshida et al., 2009). The average copy numbers of the Venus gene in the oxytocin receptor-Venus knock-in heterozygous and wild-type mice were calculated to be 1.00 and 0.00, respectively (Fig. 2A). These results indicated that the copy number of the Venus gene integrated in the mouse genome was accurately detected.

**Fig. 2.**
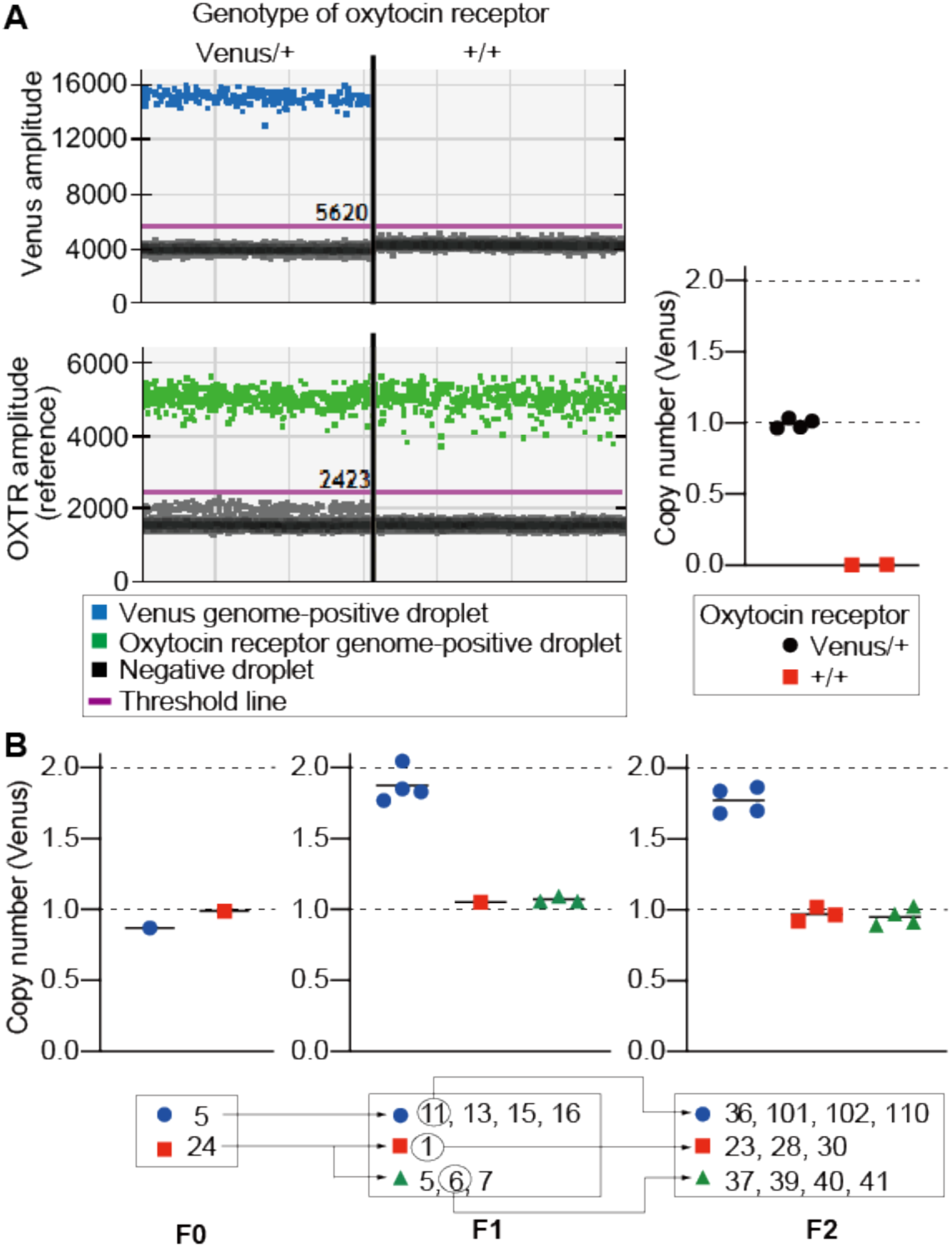
Analysis of copy number of the Venus gene by using droplet digital PCR. (A) Reliable detection of the copy number of the Venus gene by using genomic DNA of oxytocin receptor-Venus knock-in heterozygous mice generated by embryonic stem cells. Representative droplet plots of oxytocin receptor Venus/+ and +/+ (Venus-positive droplets (Upper left), OXTR positive-droplet (lower left)). Calculated copy number of the Venus gene in oxytocin receptor Venus/+ and +/+ mice (right). (B) Calculated copy numbers of the Venus gene in F0, F1 and F2 generations of Prlhr knock-in mice. Black arrows show the parent-offspring relationships.

We then conducted droplet digital PCR for two candidates of the F0 generation. The copy numbers of the Venus gene in numbers 5 and 24 of F0 were calculated to be 0.87 and 0.99, respectively (left panel of Fig. 2B). In the F1 generation, the copy number of number 1 of F1 was calculated to be 1.05 and those of numbers 5, 6, 7 of F1 were calculated to be 1.06, 1.06 and 1.10, respectively. These results indicated that the offspring of number 24 of F0 had a copy number similar to that of their parent. On the other hand, although the copy number of number 5 of F0 was 0.87, number of 11 of F1, the offspring of number 5 of F0, had approximately two copies (1.85). Numbers 13, 15 and 16, siblings of F1 number 11, also had approximately two copies (1.77, 1.83 and 2.05, respectively) (middle panel of Fig. 2B). The copy number of Venus in F2 was similar as that of their parents (right panel of Fig. 2B).

From the droplet digital PCR analysis, both of the two F0 generation mice that were candidates by conventional PCR analysis were identified as having one copy. However, analysis of their F1 and F2 generation progenies showed that F0 number 5 had two copies of the transgene and F0 number 24 had one copy insertion.

### Southern blot analysis

We performed Southern blot analysis to characterize Prlhr locus-specific targeting. Restriction enzymes BamHI and HpaI were selected from a putative designed knock-in allele for digestion of genomic DNA (Fig. 3A). 5’ and 3’ probes external to the targeting vector and an internal Venus probe were used to distinguish the wild-type (9.4-Kbp band for the 5’ and 3’ probes and no band for the Venus probe) and designed target allele (4.1-Kbp for the 5’ probe and for the Venus probe and 6.2-Kbp for the 3’ probe) (Fig. 3A). In numbers 5 and 24 of the F0 generation, the expected 4.1-Kbp band was detected by the 5’ probe (red arrowhead, left upper panel of Fig. 3B). In number 24 of F0, an unexpected weak band larger than 9.4 Kbp was also detected by the 5’ probe (green arrowhead, left upper panel of Fig. 3B). In number 5 of F0, a band larger than expected 6.2 Kbp was detected by the 3’ probe (blue arrowhead, left middle panel of Fig. 3B). In number 24 of F0, an expected 6.2-Kbp band (red arrowhead, left middle panel of Fig. 3B) and an unexpected weak band larger than 9.4 Kbp (green arrowhead, left middle panel of Fig. 3B) were detected by the 3’ probe. In number 5 of F0, an expected 4.1-Kbp band (red arrowhead, left bottom panel of Fig. 3B) and an unexpected weak band larger than 4.1 Kbp (blue arrowhead, left middle panel of Fig. 3B) were detected by the Venus probe. In number 24 of F0, an expected 4.1-Kbp band (red arrowhead, left bottom panel of Fig. 3B) and an unexpected weak band larger than 10 Kbp (blue arrowhead, left middle panel of Fig. 3B) were detected by the Venus probe. In the F1 generation, number 11 of F1 showed a pattern of bands similar to that of her parent, number 5 of F0 (middle top, middle middle and middle bottom panels of Fig. 3B). Numbers 1 and 6 of F1, offsprings of number 24 of F0, showed different band patterns. In number 1 of F1, bands of 4.1 Kbp for the 5’ probe and for the Venus probe and a band of 6.2 Kbp for the 3’ probe were detected as designed (middle top, middle middle and middle bottom panels of Fig. 3B). On the other hand, in number 6 of F1, a band larger than 9.4 Kbp for the 5’ and for 3’ probes and a band larger than 10 Kbp for the Venus probe were detected (middle top, middle middle and middle bottom panels of Fig. 3B). In the F2 generation, the detected band patterns were exactly the same as those of their parents (F1 mice) (right upper, right middle and right bottom panels of Fig. 3B).

**Fig. 3.**
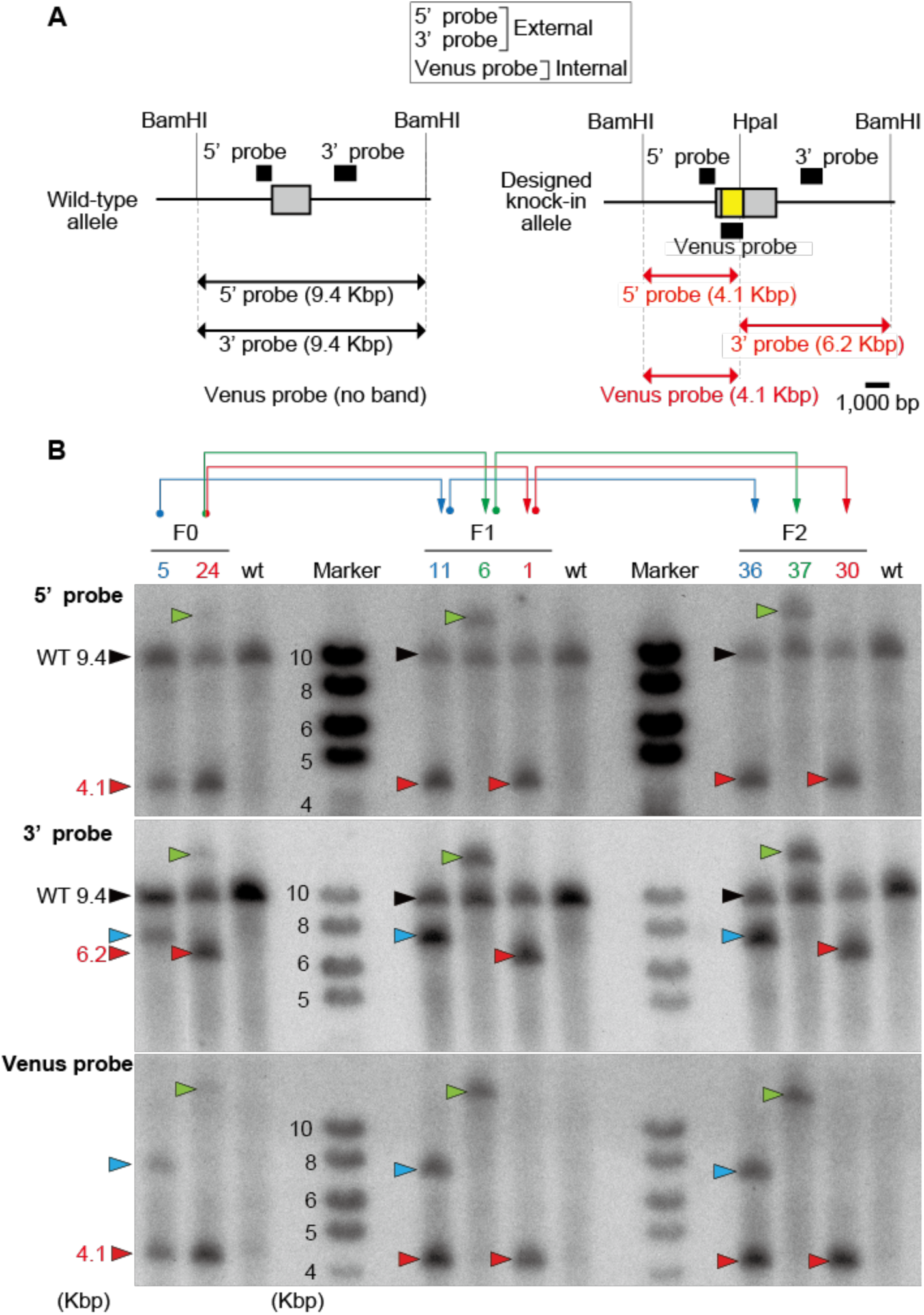
Analysis of targeted gene recombination by using of Southern blotting. (A) Schematic representation of wild-type mouse Prlhr genomic locus (left) and genomic locus with designed recombination (right). The black bars show specific probes used in Southern blotting. The horizontal arrows denote the expected sizes of restriction DNA fragments. (B) Southern blot analysis of genomic DNA of the F0, F1, F2 generations and wild type (wt). Blue, green and red arrows show the parent-offspring relationships. Black and red arrowheads show the DNA fragment from wild-type and designed knock-in allele, respectively. Green and blue arrowheads show the DNA fragments from unintentional mutant alleles.

In Southern blot analysis, none of the F0 mice showed the designed band pattern. For F0 number 5, the 5’ probe detected the expected band, but the 3’ probe detected a band that was larger in size than expected, and the Venus probe detected two bands, one expected and another unexpected. The F1 and F2 generation progenies of F0 number 5 showed exactly the same band pattern. For F0 number 24, all probes detected two bands of the mutant, one expected and another unexpected. Based on the results of conventional PCR analysis, the F1 and F2 generation progenies were divided into two types. As a result, we were able to divide the progenies into those that showed the expected band pattern for all probes (F1 number 1 and F2 number 30) and those that showed unexpected bands for all probes (F1 number 6 and F2 number 37). Unexpected bands were also detected with the 5’ and 3’ probes designed outside the targeting vector sequence, indicating that an unexpected insertion of the Venus gene occurred between the two BamHI sites in vicinity of the Prlhr locus.

### Next-generation sequencing-based RAISING analysis

We performed random integration analysis with next-generation sequencing. This method was developed for sensitive detection of clonality of cells infected with Human T-cell leukemia virus type-1, which causes adult leukemia/lymphoma (Wada et al., 2022). In number 5 of the F0 generation, two types of sequence including the Venus sequence were detected (Fig. 4A, Figure SI 1 and Table S1). Type (a) contained both the endogenous genomic sequence and the knock-in vector-containing sequence. The site of integration of Venus was at the designed location of the Prlhr gene on chromosome 19. Type (b) consisted of only the knock-in vector-containing sequence, and parts of the 3’ and 5’ arms were inverted. Among the total of 463,463 reads, the proportions of type (a) and (b) were 78.2% and 21.6%, respectively (Fig. 4B and Table S1). In number 24 of F0, type (a) was detected and the proportion of type (a) was 99.8% out of the total of 456,263 reads. In the F1 generation, results similar to those for their parents were obtained in number 11 of F1, offspring of number 5 of F0 or numbers 1 and 6 of F1, and offspring of number 24 of F0 (Fig. 4B and Table S1). Even in the F2 generation, results similar to those for their parents were obtained in number 36 of F2, offspring of number 11 of F1 or number 30 of F2, and offspring of number 1 of F1 or number 37 of F2, offspring of number 6 of F1 (Fig. 4B and Table S1).

**Fig. 4.**
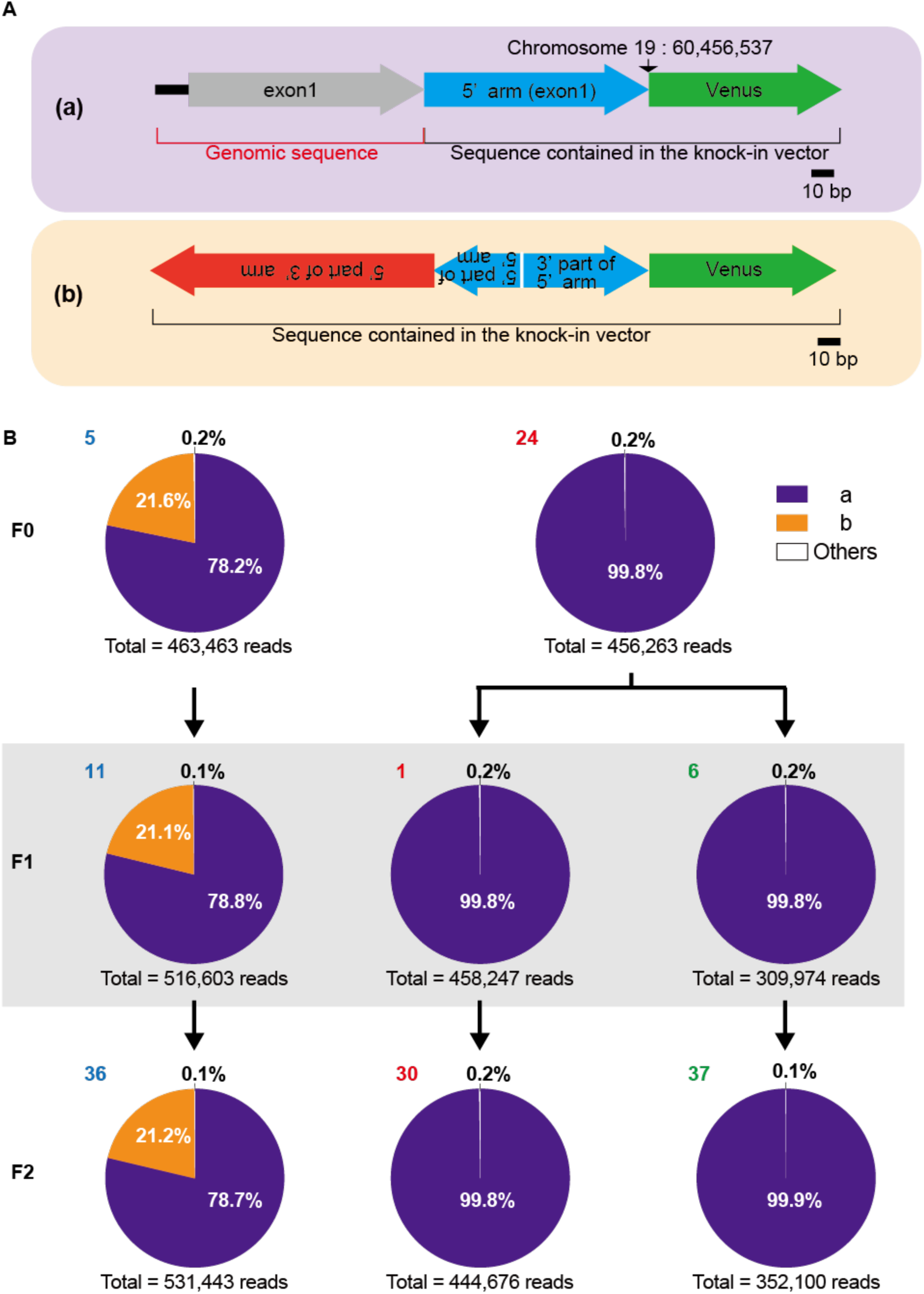
Analysis of the Venus gene integration site by using next-generation sequencing based-RAISING. (A) Schematic representation of two PCR products, type (a) and type (b), containing Venus sequences. (B) Proportion of read counts for PCR products containing Venus sequences from F0, F1, and F2 generations. Black arrows show the parent-offspring relationships.

By RAISING analysis, two different sequences including the Venus sequence were detected in F0 number 5. This result suggests that tandem two-copy occurred at one Prlhr locus or that the designed one copy knock-in occurred at one Prlhr locus and an inverted insertion of the targeting vector occurred at the other Prlhr locus. However, these two sequences did not separate in the F1 and F2 generation progenies of F0 number 5, suggesting that insertion of tandem two-copy at one Prlhr locus occurred. Although the F1 and F2 generation progenies of F0 number 24 were divided into two strains according to the results of conventional PCR analysis, only type (a) was detected in both strains. These results suggest that F1 number 6, and F2 number 37 have a Prlhr locus where Venus was inserted and genome rearrangement occurred in the vicinity.

### Genomic sequencing analysis

We performed sequence analysis of numbers 5 and 24 of the F0 generation and numbers 11 and 1 of the F1 generation. We used three primer pairs external to the targeting vector and internal primers in combination as shown in Fig. 1. Sequence analysis for PCR products of 5’ ext-1 revealed that numbers 5 and 24 of F0 and numbers 11 and 1 of F1 had accurate conjunction without deletion, insertion and mutation from upstream of exon 1 to the 5’ part of the Venus-polyA signal compared to the sequence of the designed knock-in allele (Fig. 5A, B and Figure SI 2). However, no PCR product of 5’ ext-1 was obtained for number 6 of F1, and thus we were unable to perform sequencing analysis (Fig. 1D). On PCR products of 3’ ext-1, numbers 5 and 24 of F0 and numbers 11 and 1 of F1 had exact connection and the sequence from the 3’ part of the Venus-polyA signal to the middle part of exon 1 (Fig. 5A, C and Figure SI 3). The sequence of number 6 of F1 was unable to be determined because the PCR product of the 3’ region was also not obtained (Fig. 1D). Sequence analysis for PCR products of Full ext-1 revealed that number 24 of F0 and her offspring, number 1 of F1, had whole sequences for the Venus-polyadenylation signal cassette and the sequences were exactly the same as the sequence of the designed knock-in allele (Fig. 5A, D and Figure SI 4). On the other hand, PCR products were not detected in number 5 of F0 and numbers 11 and 6 of F1 (Fig. 1D). In number 1 of F1, PCR products were also detected in conventional PCR using primer pairs located outside of the primers used for Full ext-1 as well as the primer located downstream of exon 1 and the internal primer (Figure SI 5A and B). No unexpected mutations in the confirmed range from exon 1 upstream to downstream of the knock-in allele were detected by sequence analysis of those PCR products (Figure SI 5C, D, Figure SI 6 and Figure SI 7).

**Fig. 5.**
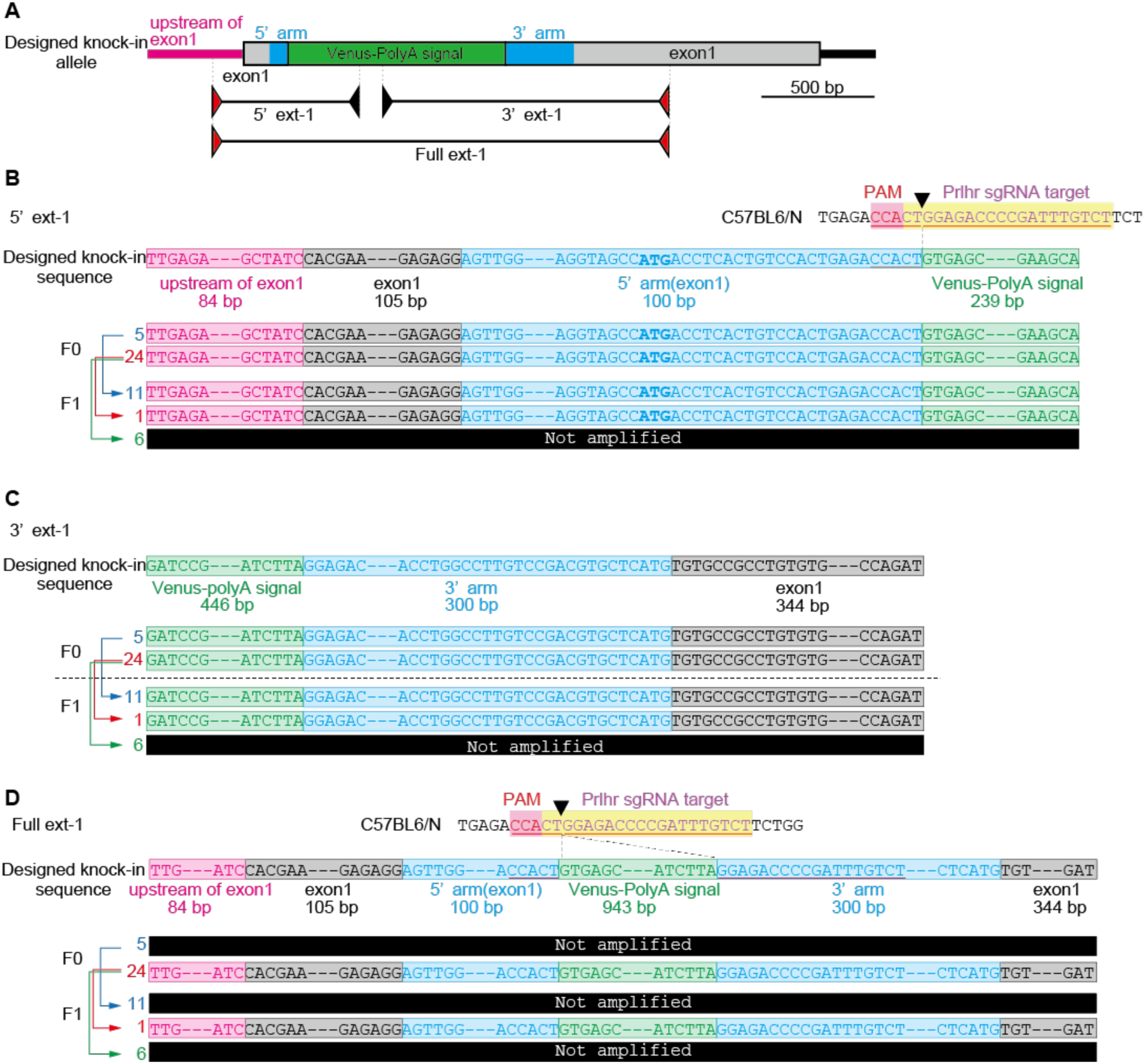
Analysis of genomic sequencing in F0 and F1 generations. (A) Schematic representation of designed Prlhr knock-in locus and primer sets for PCR amplification (primer external to the targeting vector and internal primer pairs, 5’ext-1 and 3’ext-1: primers external to the targeting vector pair, Full ext-1). Red and black arrows show the primer external to the targeting vector and internal primer, respectively. Sequence analysis for PCR products of 5’ ext-1 (B), 3’ ext-1 (C) and Full ext-1 (D). Blue, red and green arrows show the parent-offspring relationships.

In the sequence analysis, the sequences of the PCR products in the 5’ and 3’ joint areas were the designed sequences for numbers 5 and 24 of F0 and their offspring numbers 11 and 1 of F1. PCR products containing the full length of the targeting vector were analyzed, and sequences of the designed mutant alleles were detected in number 24 of F0 and number 1 of F1. These results indicated that two candidate mice of the F0 generation had Venus inserted in the Prlhr locus and that each sequence was correctly transmitted to their offsprings.

## Discussion

In this study, we conducted a detailed analysis of the generation process of a Prlhr-Venus knock-in mouse line by using five methods over three generations. After a comprehensive analysis, we found that one F0 generation mouse had the designed knock-in allele, and we were able to establish a knock-in mouse line with the designed mutant allele. Number 5 of F0 had a tandem two-copy mutant allele. F0 number 24 was a mosaic mouse with the designed mutant allele and an allele that underwent reconstruction near the prlhr locus with the targeted knock-in. These two mutant alleles were separated in the F1 and F2 generations.

### Validation of conventional PCR

In conventional PCR, the use of primers external to the targeting vector was effective because they could confirm knock-in to the target locus. On the other hand, PCR products on the 5’ and 3’ parts using the external and internal primer pairs were detected even in the two-copy mutant, suggesting that this result did not guarantee a designed one-copy mutant. The multicopy event occurred in this one guide RNA - one donor DNA method as well as in the previously reported conditional knockout method using two guide RNA - one donor DNA containing two loxP sequences (Skryabin et al., 2020). In this study, PCR products containing the full length of the target vector were detected in mice with a one-copy mutant allele using external primer pairs.

The detection of this PCR product seems to be the most reliable for detecting a one-copy mutant allele using conventional PCR. However, in a study on generation of conditional knock-out by the 2-guide RNA - one-lssDNA method, no PCR products were detected using external primer pairs despite the designed one-copy integration (Skryabin et al., 2020). In addition, in the majority of studies on improved knock-in methods using CRISPR-cas9, it was not determined whether PCR products were detected or not by using external primer pairs. (Gu et al., 2018; Lin et al., 2018; Ma et al., 2014; Yao et al., 2017; Yao et al., 2018; Yoshimi et al., 2016; Yoshimi et al., 2021), although the product was reported in one paper (Quadros et al., 2017). PCR amplification with external primer pairs is important for confirmation of a single copy number of the transgene, although the capability of PCR amplification may depend on the characteristics of the targeting vector and the target locus and thus the fact that the product is not detected does not necessarily mean that the expected mutation did not occur.

Number 24 of F0 was a mosaic genotype with two different knock-in mutant Prlhr alleles. This mosaicism was not able to be detected by conventional PCR using the genome of number 24 of F0 and could only be detected by using multiple primer pairs for the genomes of F1 and F2 generations. Conventional PCR is most often used for genotyping for strain maintenance. At least for the F1 generation, multiple external primer pairs should be used to select rodents with the designed mutation and to remove unexpected mutants.

### Validation of droplet digital PCR

We first confirmed the accuracy of copy number detection by droplet digital PCR using oxytocin receptor-Venus knock-in heterozygous mice established with ES cells via homologous recombination. In the copy number analysis of the Venus gene using these mice, we were able to detect the one-copy mutant with high confidence. In Prlhr-Venus knock-in mice, number 5 of the F0 generation was identified as a one-copy mutant. On the other hand, all heterozygous progenies of number 5 of F0 were detected as two-copy mutants over the F2 generation. HDR-mediated repair occurs during the S and G2 phases, when sister chromatids are prepared (Gilbert, 2006). There can therefore be temporarily four Prlhr loci in a one-cell fertilized egg. In the case of number 5 of F0, the two targeting vectors were inserted into one locus in tandem, and thus when droplet digital PCR was performed on the genomic DNA extracted from the F0 mouse, it was detected as one copy (Fig. 6A). Number 24 of F0 was identified as a one-copy mutant and all heterozygous progenies of the mouse were also detected as one-copy mutants. In this case, one targeting vector was inserted into two of the four loci, and therefore the results of droplet digital PCR were calculated as one copy (Fig. 6B).

**Fig. 6.**
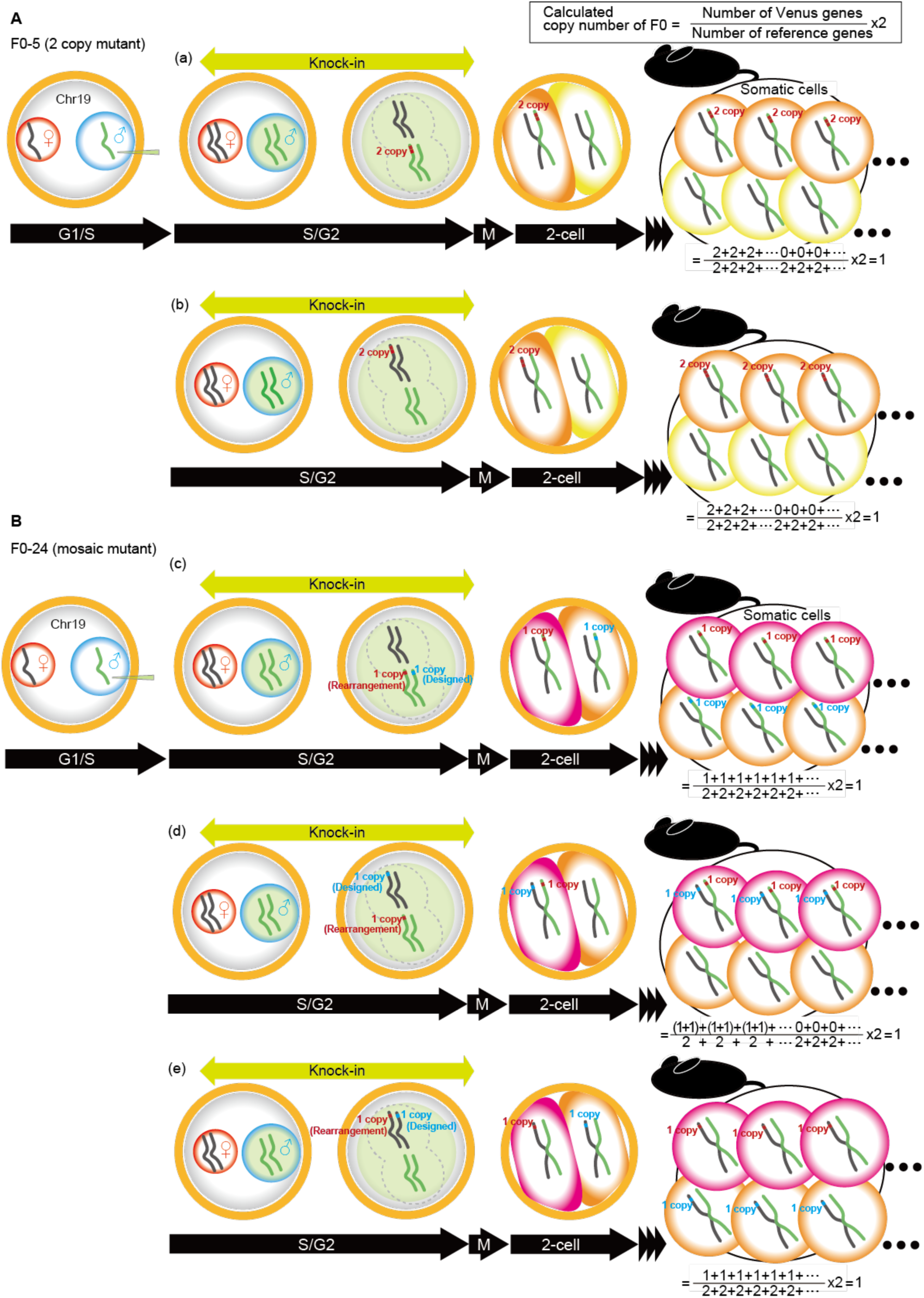
Schematic drawing of the estimated knock-in events that occurred at the Prlhr locus in the F0 generation. A cocktail of hCas9 mRNA, sgRNA and lssDNA was injected into the male pronuclei of embryos. HDR-mediated knock-in events occur in the S/G2 phase of the pronuclear stage. (A) In F0 number 5, tandem two-copy insertions occurred on (a) one paternal or (b) one maternal chromosome. In droplet digital PCR using the genome extracted from the tail of F0 number 5, the copy number was calculated to be a single copy mutant in either case. (B) In F0 number 24, single-copy insertions occurred in two different Prlhr loci, and one KI allele was as designed and the other was an allele in which rearrangement occurred: (c) two paternal chromosomes, (d) one paternal chromosome and one maternal chromosome, or (e) two maternal chromosomes. In droplet digital PCR using the genome extracted from the tail of F0 number 24, the copy number was also calculated to be a single copy mutant in all cases.

For F1 and F2 heterozygous mice, the copy number obtained by droplet digital PCR is quite reliable because the knock-in allele and the wild-type allele are present in a one-to-one ratio (Figure SI 8). In F0 mice, there is a possibility that multi-copy or mosaic genotypes exist, and the copy number cannot be calculated exactly. However, the estimation of knock-in events occurring in F0 embryos by comparing results of copy numbers in F0 and F1 with droplet digital PCR is valuable information for the development of efficient and appropriate knock-in methods.

### Validation of Southern blotting

Although Southern blotting is a classical method, we were able to detect both designed and unexpected mutant loci. In order to obtain sufficient information from the genomic DNA of the F0 generation, it was necessary to use all 5’ and 3’ probes external to the targeting vector and internal Venus probe.

In number 5 of F0, the band pattern was detected as expected in the 5’ probe. However, the band detected with the 3’ probe showed a larger size than expected, and the Venus probe also detected a band other than the expected one. These results suggested that knock-in occurred at the Prlhr locus but with unexpected multicopy mutation, which was unable to be detected by the 5’ probe alone. For the F0 number 24, all probes detected unexpected bands in addition to the expected bands. This suggests that the mice had two different mutations at the Prlhr loci. According to the PCR results, the progenies were divided into two genotypes, and it was found that F0 had a mosaic of two knock-in genotypes. Because a larger band than that of the wild type was detected by the external probes, rearrangement occurred in the vicinity of the prlhr locus.

External probes can detect gross genomic changes near the target locus, while internal probes can detect integration events including off-target insertions from the entire genome. However, the weakness of Southern blotting is that it cannot detect small insertion or deletion mutations around the target locus, which are often observed in the CRISPR-Cas9 method (Yoshimi et al., 2016; Yoshimi et al., 2021), if they occur at the same time as the knock-in event to the target locus.

### Validation of next-generation sequencing-based RAISING

The RAISING method was used to detect knock-in in the target locus and off-target insertions in this study. Two different sequences were detected for number 5 of the F0 generation. Type (a) contained sequences outside the targeting vector and showed the designed knock-in. Type (b) contained sequences with partially inverted 5’ and 3’ homology arms but did not include genomic sequences outside the targeting vector. This result was expected to result in either a tandem two-copy at one Prlhr locus or a designed one-copy knock-in at one Prlhr locus and a reverse insertion of the targeting vector at the other Prlhr locus. The results of Southern blotting of F0 number 5 showed that the designed knock-in-derived band was not obtained with the 3’ probe, suggesting that a tandem two-copy form likely occurred. Both types of sequences were also detected in the progeny of number 5 of F0, and their proportions of sequence reads were the same as in F0. The fact that type (a) and type (b) did not separate in F1 and F2, that the sequence of type (b) contained the sequence of the 3’ arm region and that all heterozygous progenies of number 5 of F0 were also detected as two-copy mutants in droplet digital PCR analysis suggested that the sequences containing these two Venus genes were arranged in tandem at the Prlhr locus. In number 24 of F0, only the type (a) sequence was detected. According to the results of conventional PCR, the strain was divided into two groups, but only type (a) was detected in all F1 and F2 mice. Number 6 of F1 and number 37 of F2 were predicted to have off-target insertions because no PCR products were detected by conventional PCR using external primers. However, combined with the results of Southern blotting, in which the mutant allele was detected with the 5’ and 3’ probes, targeted knock-in was found at the Prlhr locus but with reconstruction occurring in the vicinity. Unlike conventional PCR, for which results depend on the selection of primer pairs, the RAISING method provides information on the insertion of exogenous gene sequences from the entire genome at the sequencing level. However, the weakness is that the maximum sequence length per read is 300 bp, and unless the homologous arm sequence is less than 250 bp, the sequence cannot reach the endogenous genomic sequence and the insertion position cannot be identified.

### Validation of genomic sequencing

Sequence confirmation by sequencing is important for the CRISPR-Cas9 method, which can induce insertion and deletion mutations. These mutations can occur in the junction region along with the knock-in of the targeting vector (Yoshimi et al., 2016; Yoshimi et al., 2021). In the PCR products of the 5’ and 3’ portions using external primers, number 5 of the F0 generation and its offspring, number 11 of F1, showed all the designed junction sequences. These results indicate that correct junctions were generated at the 5’ and 3’ parts in tandem multicopy mutants and that even if the sequences of the 5’ and 3’ junctions are correct, they cannot guarantee a single copy mutant. In number 24 of F0, the PCR product containing the full length of the targeting vector using external primers was confirmed to be the designed sequence. In this study, lssDNA was used as donor DNA, and no unexpected mutations were found in all of the sequences analyzed, including the Venus - polyA signal cassette. However, in a previous study using two gRNAs and one lssDNA to generate conditional knock-out mice, unexpected point mutations occurred in the sequence within the targeting vector (Codner et al., 2018). It is necessary to verify the sequencing within the targeting vector as well as the sequencing at the 5’ and 3’ junctions.

### Four possible events in knock-in by the CRISPR-cas9 method

In the generation of knock-in mice by homologous recombination of ES cells, chimeric mice can be obtained by using a single clone with the correct recombination selected from a large number of ES cell clones. Therefore, chimeric mice, mice in the F0 generation, have a unitary knock-in allele. The knock-in allele of the F1 mice obtained by crossing chimeric mice with the wild type is also identical to that of the injected ES cells. However, careful analysis was required for the F0 mice obtained by the CRISPR-cas9-based knock-in method, because the somatic genomic DNA was a mixture of designed and unexpected mutations. In order to identify promising F0 generations and establish designed strains, we used five genome analysis methods and detected knock-in at the target locus, genomic rearrangements in the vicinity of the knock-in allele, and multi-copy and mosaic genotypes (Fig. 7). For detection of knock-in in the target locus, conventional PCR and Southern blot analysis provided information, and the RAISING method and genome sequencing were particularly informative because they provided sequence data. This was similar for both the F0 generation and the F1 and F2 generations. Genomic rearrangements occurring in the vicinity of the knock-in allele could only be detected by Southern blotting, and this was similar for the F0 generation and the F1 and F2 generations. For detection of a multicopy, Southern blotting and the RAISING method were effective in the F0 generation, while in the F1 and F2 generations, in addition to these methods, information from droplet digital PCR with higher accuracy was useful. Mosaicism was detected by Southern blotting in the F0 generation. In the F1 and F2 generations, mosaicism was eliminated and consequently could also be identified by conventional PCR.

**Fig. 7.**
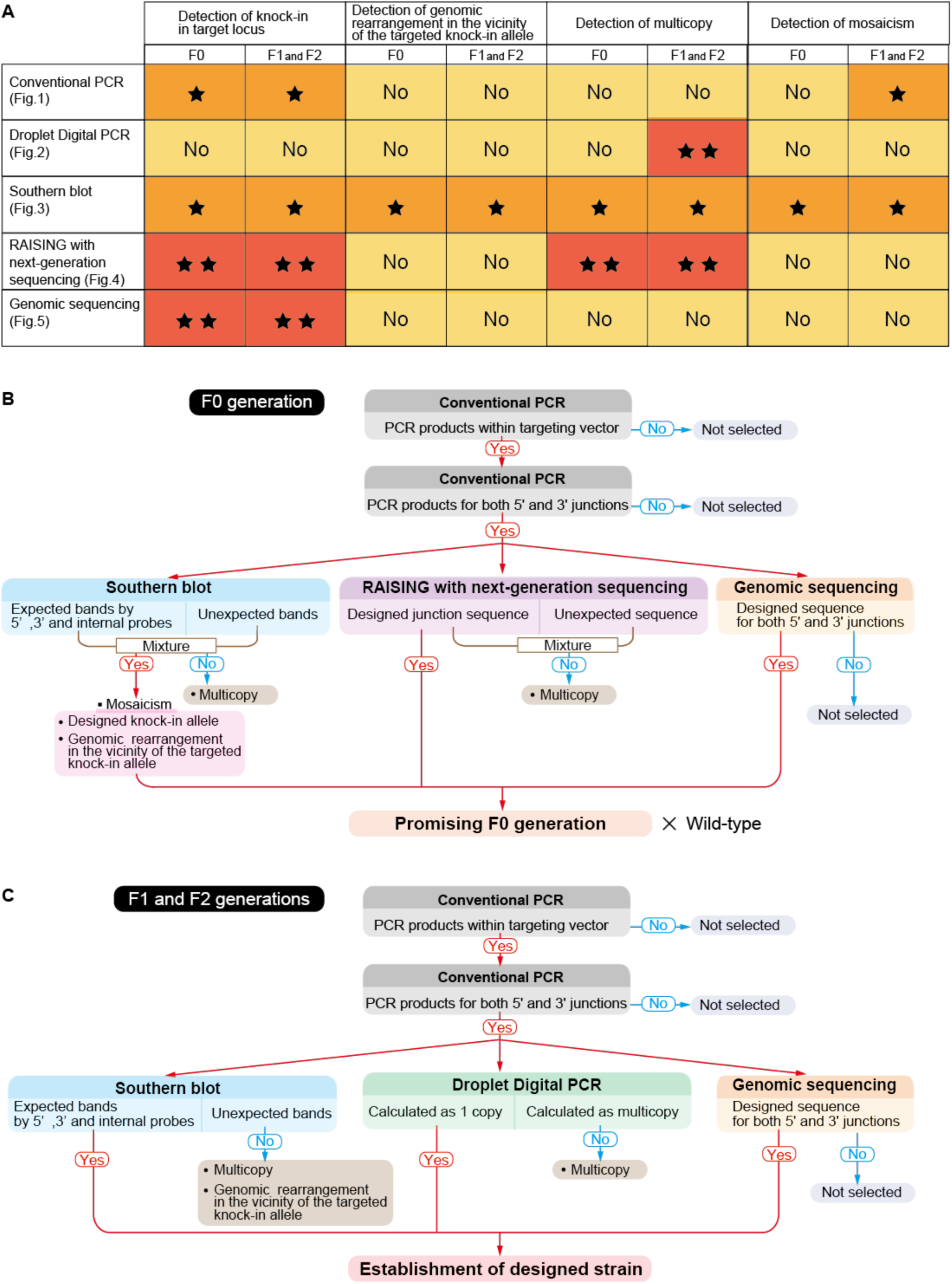
Comparison of the detection performances of five methods for four possible events in knock-in by the CRISPR-cas9 method and the process for selection of promising F0 generation and designed strain in this study. (A) According to the results obtained from this study, the superiority of each method for detecting the events was summarized. ** More informative, * Informative and ^No^ No or less information. (B) The flow of a promising F0 generation sorting process is illustrated. Southern blotting was able to detect mosaicism and multicopy. The RAISING method was capable of detecting multicopy. Conventional PCR and genome sequencing were unable to detect mosaicism and multicopy. (C) The flow for establishing a designed strain is illustrated. Southern blotting was able to identify multicopy and genomic rearrangements near the target knock-in allele. Droplet digital PCR was able to distinguish multicopy. Conventional PCR and genome sequencing were unable to detect multicopy.

## Conclusion

It was better to make a selection based on the assumption that the F0 generation mice obtained by the CRISPR-cas9-based knock-in method had a mixture of designed and unexpected mutations. Analytical methods that can revealed unexpected mutations as well as designed mutations enabled selection of promising F0 mice with confidence. Southern blotting and the RAISING method were particularly useful for detecting unexpected mutations in the whole genome. For the F0 generation, the combination of four methods, conventional PCR, RAISING, southern blotting and genome sequencing, was effective, and for the F1 and F2 generations, the results of droplet digital PCR analysis in addition to these methods were beneficial for establishing the strain. On the other hand, in the case of a tandem multicopy, regardless of the generation, conventional PCR, Southern blotting and sequencing yielded results that were partially identical to those of one-copy. These results indicate that focusing only on positive evidence can lead to erroneous selection of undesigned strains. Further development of the knock-in method using CRISPR-cas9 is expected in the future. However, the selection of F0 mice is essential for the establishment of strains with the designed genetic mutations. In this study, we investigated the effectiveness of a methodology for selecting strains with genetic mutations obtained by the knock-in method using CRISPR-cas9. This methodology is useful for accurate selection of the designed strains. In addition, being able to decide whether or not to plan to acquire F0 mice through additional experiments without waiting for the results of mutation analysis of the F1 generation can greatly contribute to saving time and effort for the experimenters.

## Materials and methods

### Animals

Mice of the C57BL/6N strain (Charles River Laboratories, Kanagawa, Japan), oxytocin receptor-Venus knock-in heterozygous mice backcrossed with C57BL/6J mice for over 18 generations (Yoshida et al., 2009), were used in the present study. The mice were housed under a 12:12- hour light/dark photocycle (lights on 7:30 AM) at 20-24°C and 40%-70% relative humidity. Food and water were available ad lib. All animal procedures were approved by the Institutional Animal Experiment Committee of Jichi Medical University and Institute of Immunonogy Co., Ltd. and were conducted in accordance with the Institutional Regulations for Animal Experiments and Fundamental Guidelines for Proper Conduct of Animal Experiments and Related Activities in Academic Research Institutions under the jurisdiction of the Ministry of Education, Culture, Sports, Science and Technology.

### Preparation of Cas9 mRNA and sgRNA

Expression vectors of hCas9 containing the T7 promoter, SV40 nuclear localization signal-fused hCas9 and 81-bp polyadenylation signal were used (Yoshimi et al., 2016). Cas9 mRNA was transcribed *in vitro* from a linearized plasmid, added poly(A) tail and purified by using the MessageMAX T7 ARCA Capped Message Transcription Kit (Cellscript, LLC., Wisconsin, USA), poly(A) Polymerase Tailing Kit (Epicentre Biotechnologies, Wisconsin, USA) and MEGAClear Kit (Thermo Fisher scientific, Inc., Massachusetts, USA). Software tools for sgRNA design, CRISPR design (Ran et al., 2013), and CRISPR direct (RRID:SCR_018186) were used for prediction of unique target sites throughout the genome sequence of the mouse. Synthesis and purification of sgRNA were performed by FASMAC Co., Ltd (Kanagawa, Japan).

### Preparation of lssDNA

A DNA fragment containing a Venus-SV40 polyadenylation signal cassette and two homology arms was cloned into pUC plasmid. A 100-bp fragment was used as the 5’ homology arm and a 300-bp fragment was used as the 3’ homology arm (Table S2). The target sequence in the plasmid was amplified using PCR with a primer pair containing a nuclease resistant primer, and the PCR product was digested with 5’-3’ exonuclease to produce lssDNA. After digesting the template plasmid, the lssDNA was sequenced and confirmed to be lssDNA by capillary electrophoresis. The lssDNA was stored at -80°C until use.

### Microinjections into mouse embryos

Female mice were superovulated by injection of pregnant mare serum gonadotropin (PMSG, ASKA Pharmaceutical Holdings Co., Ltd., Tokyo. Japan) and human chorionic gonadotropin (hCG, ASKA Pharmaceutical Holdings Co., Ltd.). Pronuclear-stage embryos were then collected from superovulated females. The embryos were cultured in a KSOM medium (ARK Resource, Kumamoto, Japan) before and after microinjections. A mixture of 200 ng/μL Cas9 mRNA, 100 ng/μL sgRNA and 50 ng/μL lssDNA was microinjected into the male pronuclei of embryos by using a micromanipulator (Narishige, Tokyo, Japan). The embryos were cultured in the KSOM medium and then transferred into pseudopregnant female mice.

### Conventional PCR and sequencing analysis for detection of genomic mutations

Genomic DNA extracted from the tail was used. Genomic PCR was performed in a 25-μL reaction volume containing HotStarTaq DNA polymerase (Qiagen, Hilden, Germany) or Q5 High-Fidelity DNA polymerase (New England Biolabs), genomic DNA and 12.5 pmol of each primer. The primers used are shown in Table S3. PCR products were directly sequenced using the BigDye terminator v3.1 and the Applied Biosystems 3130xl DNA Sequencer (Thermo Fisher Scientific, Inc.) according to the standard protocol of the manufacturer.

### Southern blot analysis for detection of mutants

Five μg of genomic DNA was digested with BamHI and HpaI (New England Biolabs, Massachusetts, USA) and loaded on 0.8% agasose gels. The digested DNA samples were subjected to electrophoresis and transferred to Hybond-XL membranes (Cytiva, Tokyo, Japan). The membranes were hybridized to ^32^P-labeled DNA probes. The probes were obtained by digestion with restriction enzymes and labeled with DNA polymerase I, Large (Klenow) Fragment (New England Biolabs) and random primers (Takara Bio Inc., Shiga, Japan) with [^32^P]dCTP (PerkinElmer, Massachusetts, USA). The probes used are shown in Table S4.

### Droplet digital PCR for determination of transgene copy number

Droplet digital PCR was performed by using a QX200 droplet digital PCR system (Bio-Rad Laboratories, Inc., California, USA). Genomic DNA was digested with the restriction enzyme TaqαI (New England Biolabs). The mouse oxytocin receptor gene was used as a reference gene for normalization of the Venus copy number. The assay was performed in a 20-μL reaction volume containing ddPCR supermix for probes (Bio-Rad Laboratories, Inc.), gene-specific primers and hydrolysis probes. Each reaction was performed in duplicate. The hydrolysis probe sets used are shown in Table S3. The hydrolysis probe set for the mouse oxytocin receptor gene was designed in exon 4. Oxytocin receptor-Venus knock-in heterozygous mice generated by embryonic stem cells were used for positive controls. In the mice, part of exon 3 was replaced with a Venus-polyadenylated signal cassette, but exon 4 was intact. Droplet digital PCR data were analyzed with QuantaSoft version 1.7 software (Bio-Rad Laboratories, Inc.), and the copy number was calculated.

### Next-generation sequencing

Rapid amplification of integration sites was performed according to a previous report with minor modifications (Wada et al., 2022). Specific primers used for the amplification in this study are shown in Table S3. The final PCR products were purified by using an Agencourt AMPure XP kit (Beckman Coulter, California, USA) and were quantified by using a Qubit dsDNA HS assay kit (Thermo Fisher Scientific, Inc.) and an Agilent BioAnalyzer with High-Sensitivity DNA chip (Agilent Technologies, California, USA). Next-generation sequencing was performed by using a MiSeq Reagent Kit v3 (600-cycle) on the Illumina MiSeq system (Illumina, California, USA) according to the manufacturer’s protocols.

For data analysis, amplicon-sequence reads less than 50 nucleotides and low-quality sequencing reads were excluded by using fastp software (Chen et al., 2018) (Table S5). Adapter sequences in the sequencing reads were also trimmed with fastp (RRID:SCR_016962). A homology search was then performed by using Magic-BLAST (RRID:SCR_015513), and trimmed sequence reads that had a sequence both of 20 or more nucleotides of Venus and 90% or more match identity with the *Mus musculus* genome were extracted. Genomic locations of Venus were determined on the *Mus musculus* genome GRCm39 for all extracted sequence reads. The extracted sequence reads were grouped on the basis of Venus insertion sites and were analyzed with SnpEff software (RRID:SCR_005191) which annotates functional effect prediction (Table S1).

## Acknowledgements

We thank Naoki Kumagai, Megumi Hata, Makiko Nakamura, Yukiko Kumakura, Shinobu Iwakiri and Yuko Sawano for technical assistance. We also thank Kayoko Koeda for secretarial assistance.

## Author contributions

M.Y., Y.Totsuka and T.O. designed research. M.Y. and T.S. performed research. M.Y., T.S. and Y.Takayanagi analyzed data. M.Y. wrote original draft. All authors edited the manuscript.

## Data availability

The data supporting the findings of this study are available from the corresponding author on request.

---

**Figure SI 1.**
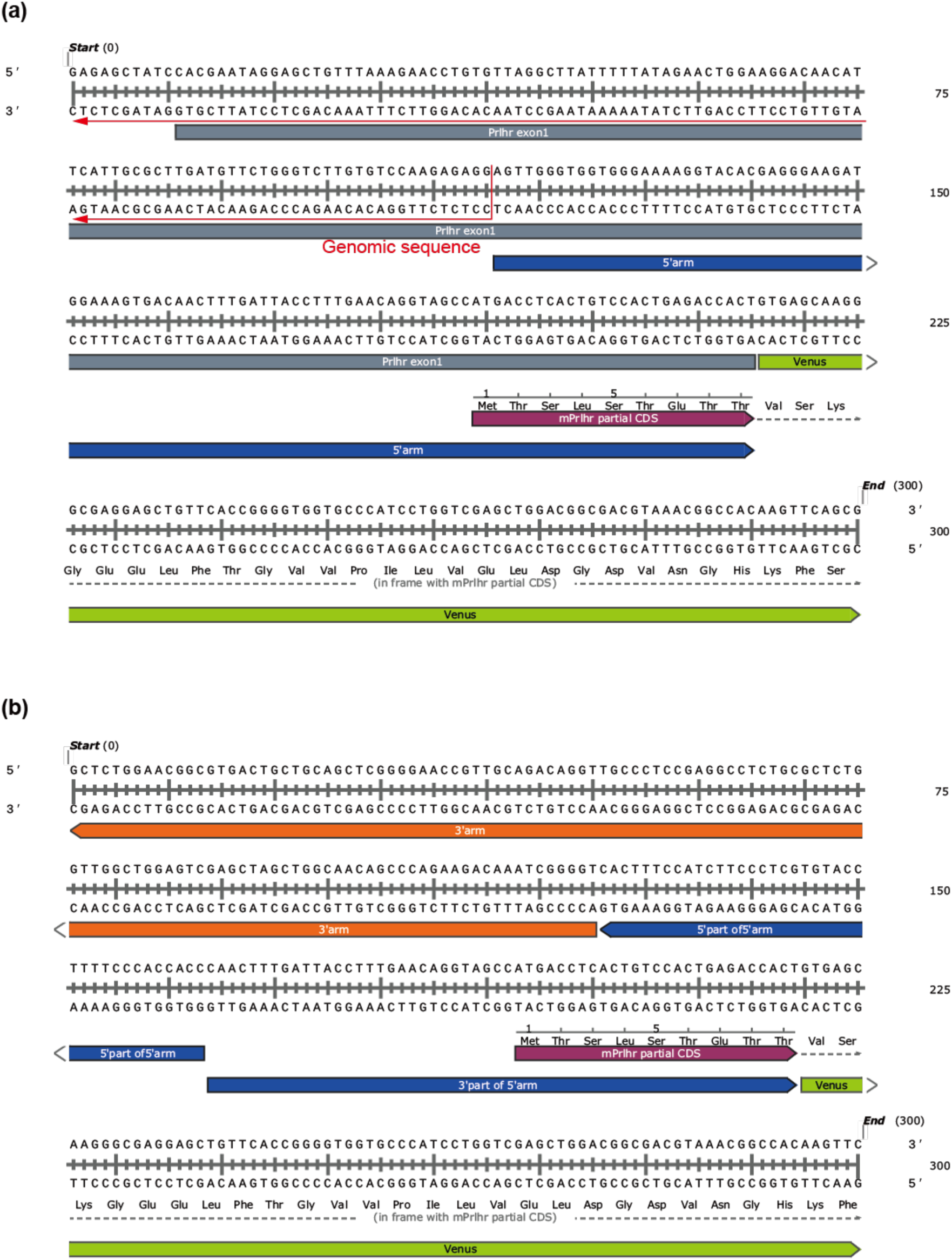
Two types of sequences including the Venus sequence detected in RAIGING analysis. Two PCR products, type (a) and type (b), containing Venus sequences are shown.

**Figure SI 2.**
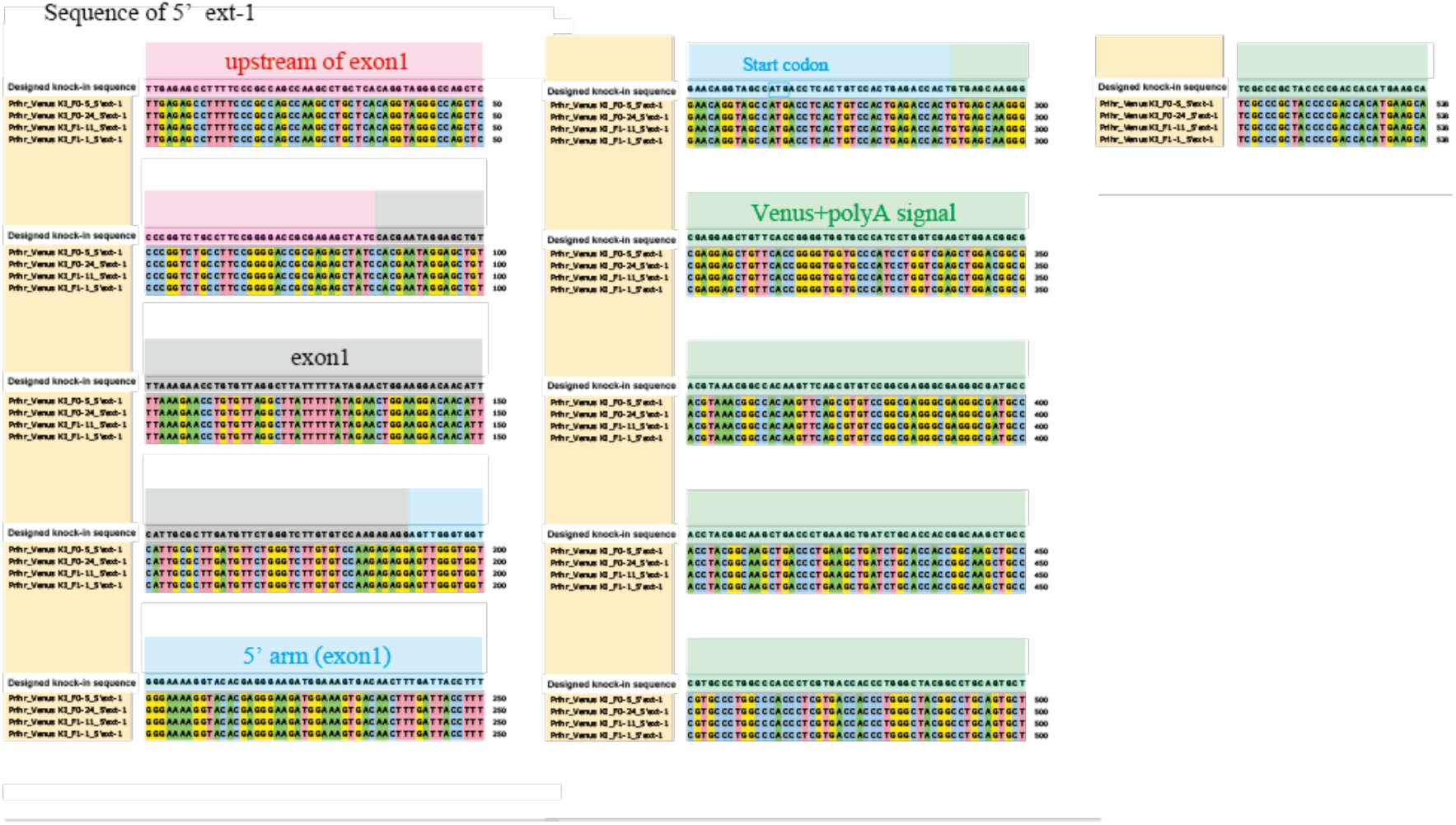
Results of mapping with Magic-BLAST and annotation with SnpEff.

**Figure SI 3.**
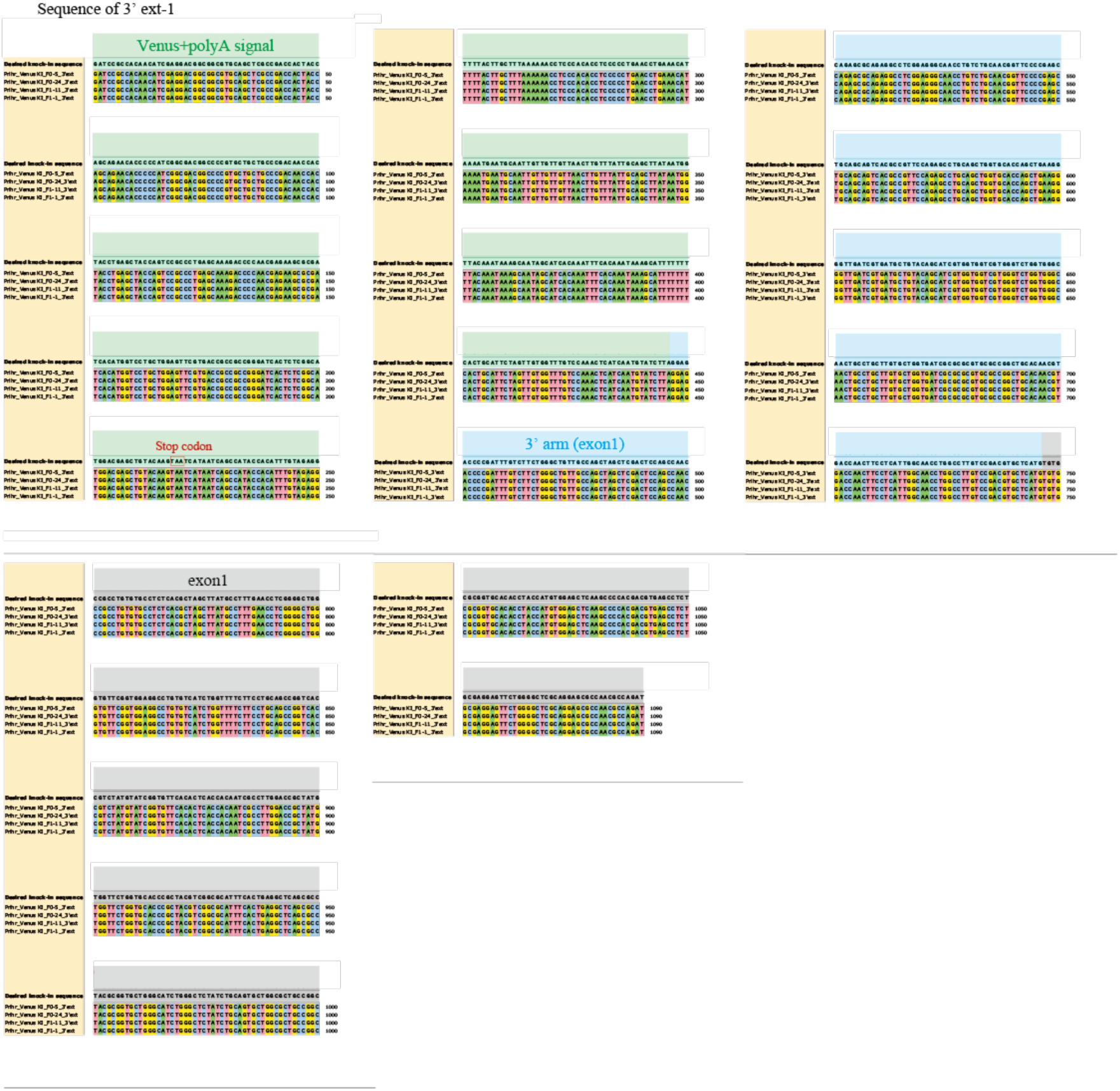
Comparison of 5’ ext-1 PCR product of the target genomic region with the designed knock-in sequence. The designed knock-in sequence was compared to the PCR product 5’ ext-1 amplified from numbers 5 and 24 of the F0 generation and numbers 11 and 1 of the F1 generation. No unexpected mutations were detected in all PCR products.

**Figure SI 4.**
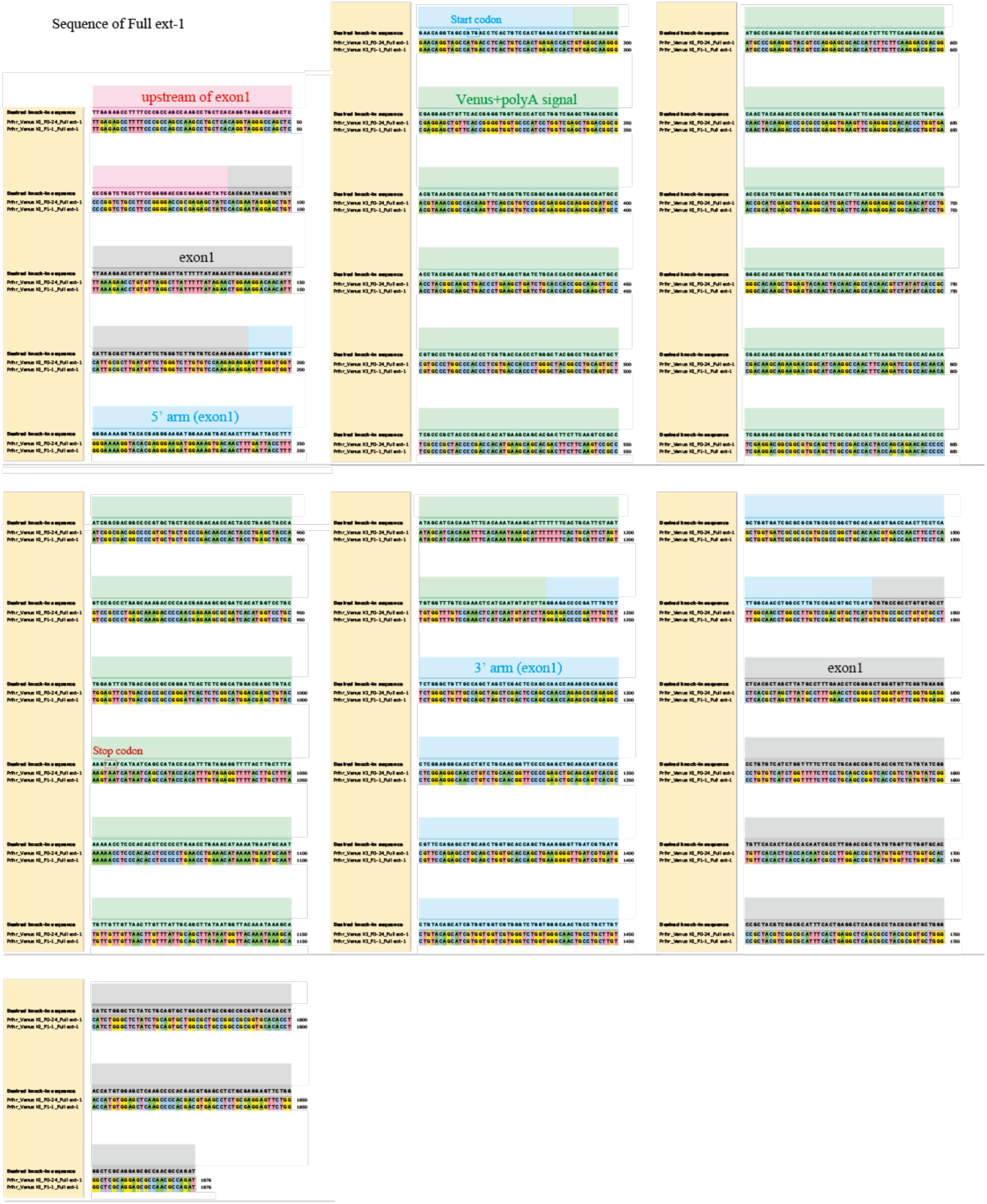
Comparison of 3’ext-1 PCR product of the target genomic region with the designed knock-in sequence. The designed knock-in sequence was compared to the PCR product 3’ ext-1 amplified from numbers 5 and 24 of the F0 generation and numbers 11 and 1 of the F1 generation. No unexpected mutations were detected in all PCR products.

**Figure SI 5.**
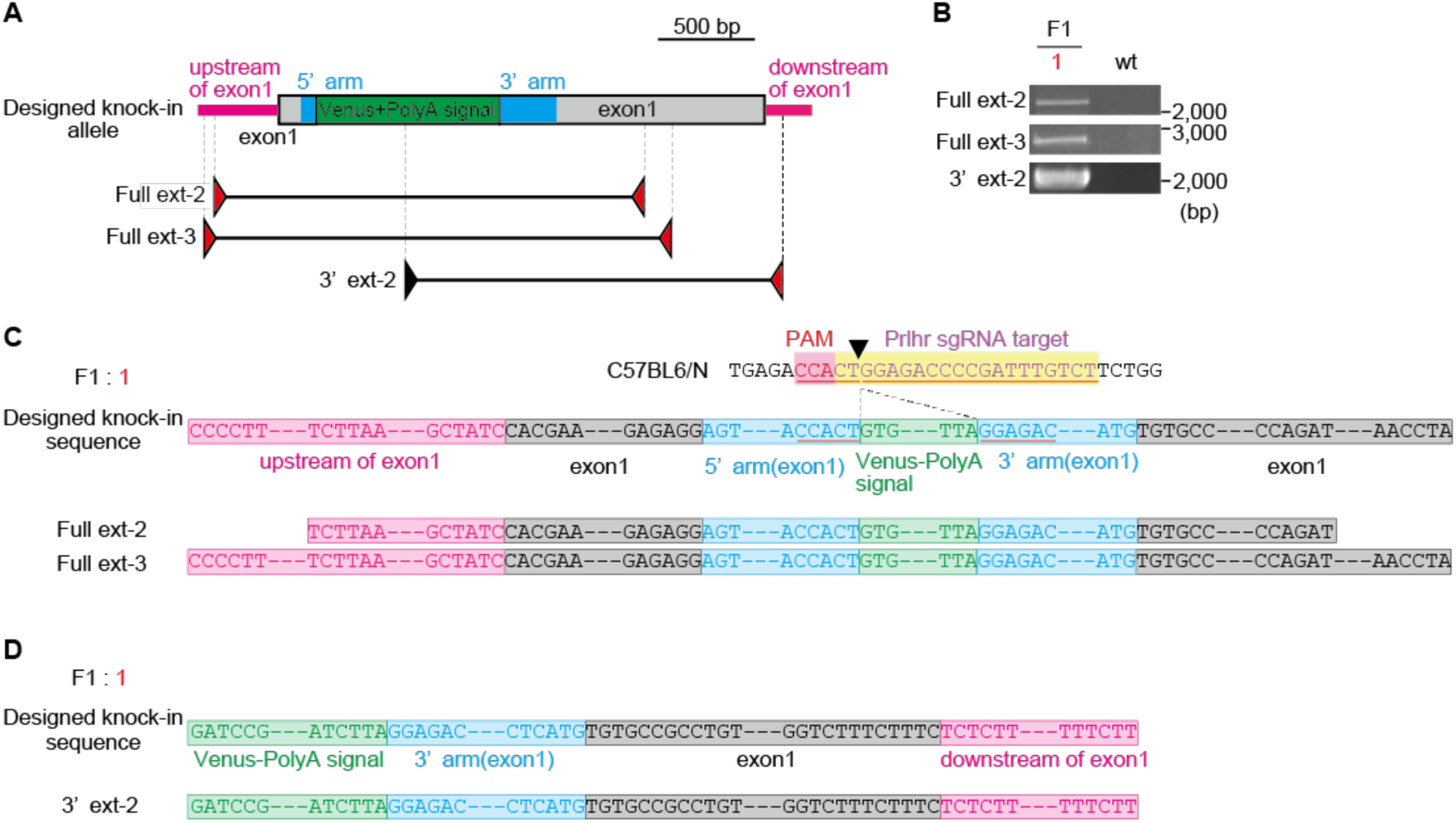
Comparison of Full ext-1 PCR product of the target genomic region with the designed knock-in sequence. The designed knock-in sequence was compared to the PCR product Full ext-1 amplified from number 24 of the F0 generation and number 1 of the F1 generation. No unexpected mutations were detected in all PCR products.

**Figure SI 6.**
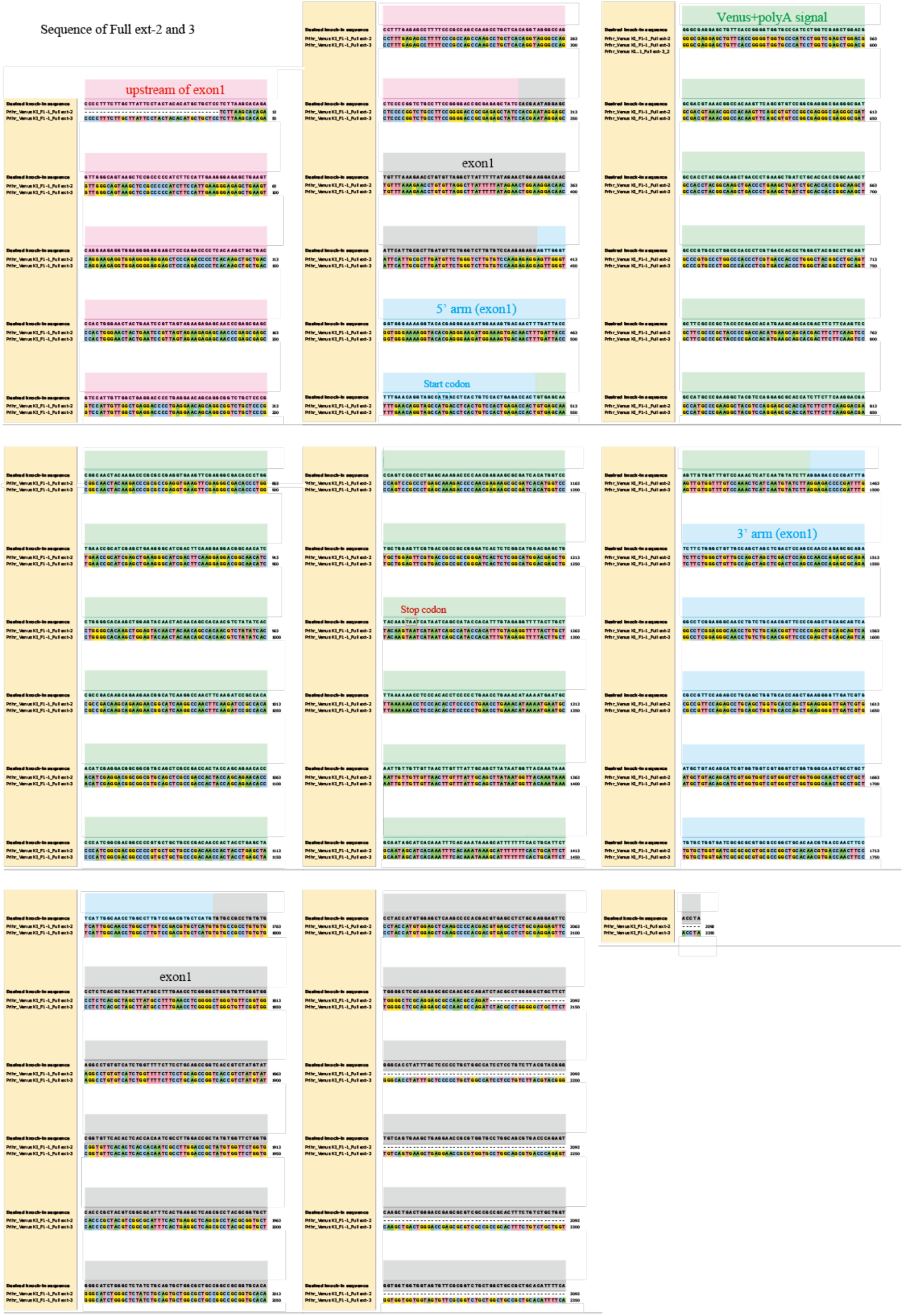
Analysis of conventional PCR and genomic sequencing in number 1 of the F1 generation. (A) Schematic representation of designed Prlhr knock-in locus and primer sets for PCR amplification (primer external to the targeting vector pairs, Full ext-2 and -3: internal primer and primer external to the targeting vector pair, 3’ ext-2). Red and black arrows show the primer external to the targeting vector and internal primer, respectively. (B) Conventional PCR analysis of genomic DNA from number 1 of the F1 generation by using each primer pair. Sequence analysis for PCR products of Full ext-2 and -3 (C) and 3’ ext-2 (D).

**Figure SI 7.**
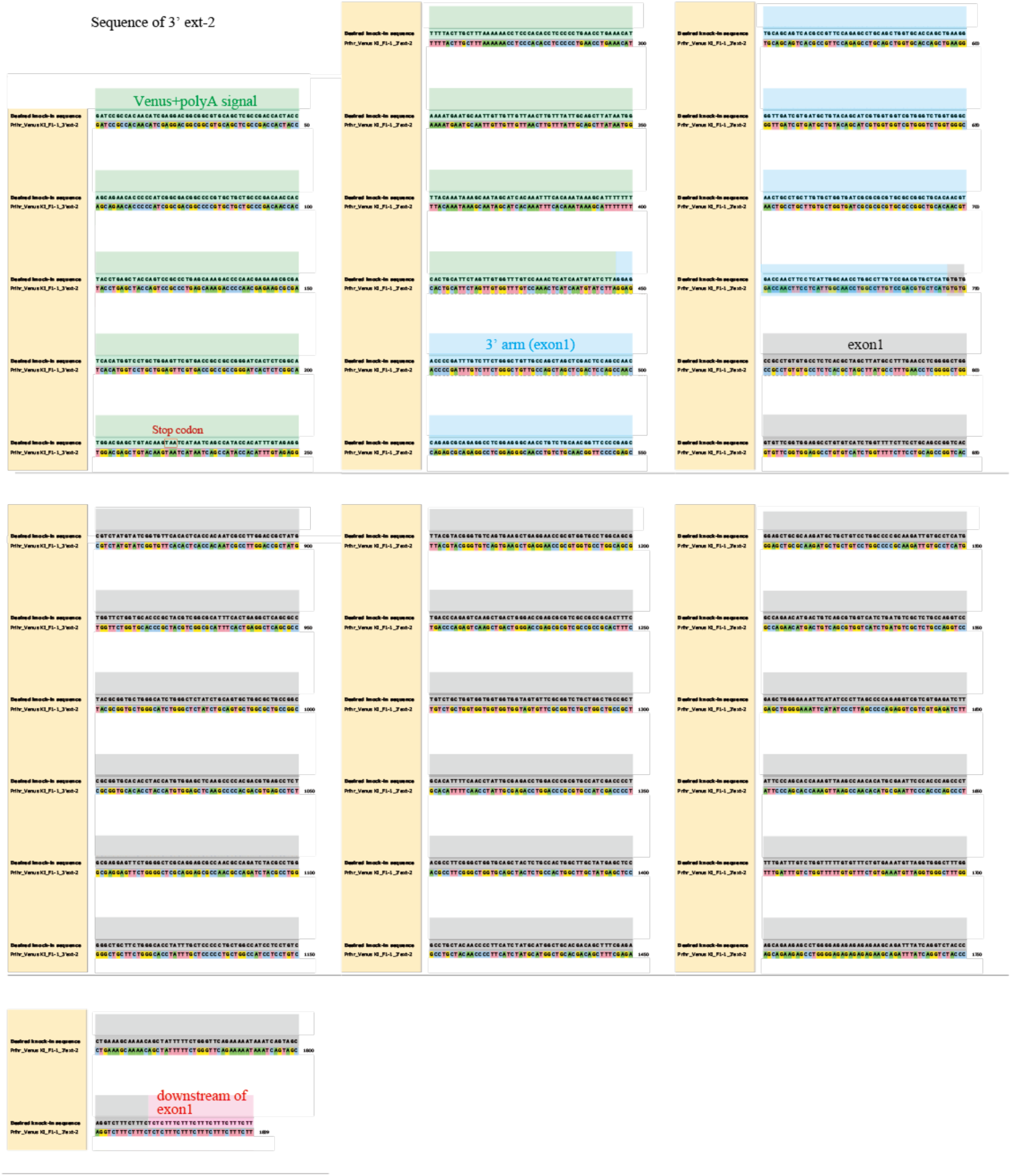
Comparison of Full ext-2 and -3 PCR products of the target genomic region with the designed knock-in sequence. The designed knock-in sequence was compared to the PCR products Full ext-2 and -3 amplified from number 1 of the F1 generation. No unexpected mutations were detected in the two PCR products.

**Figure SI 8.**
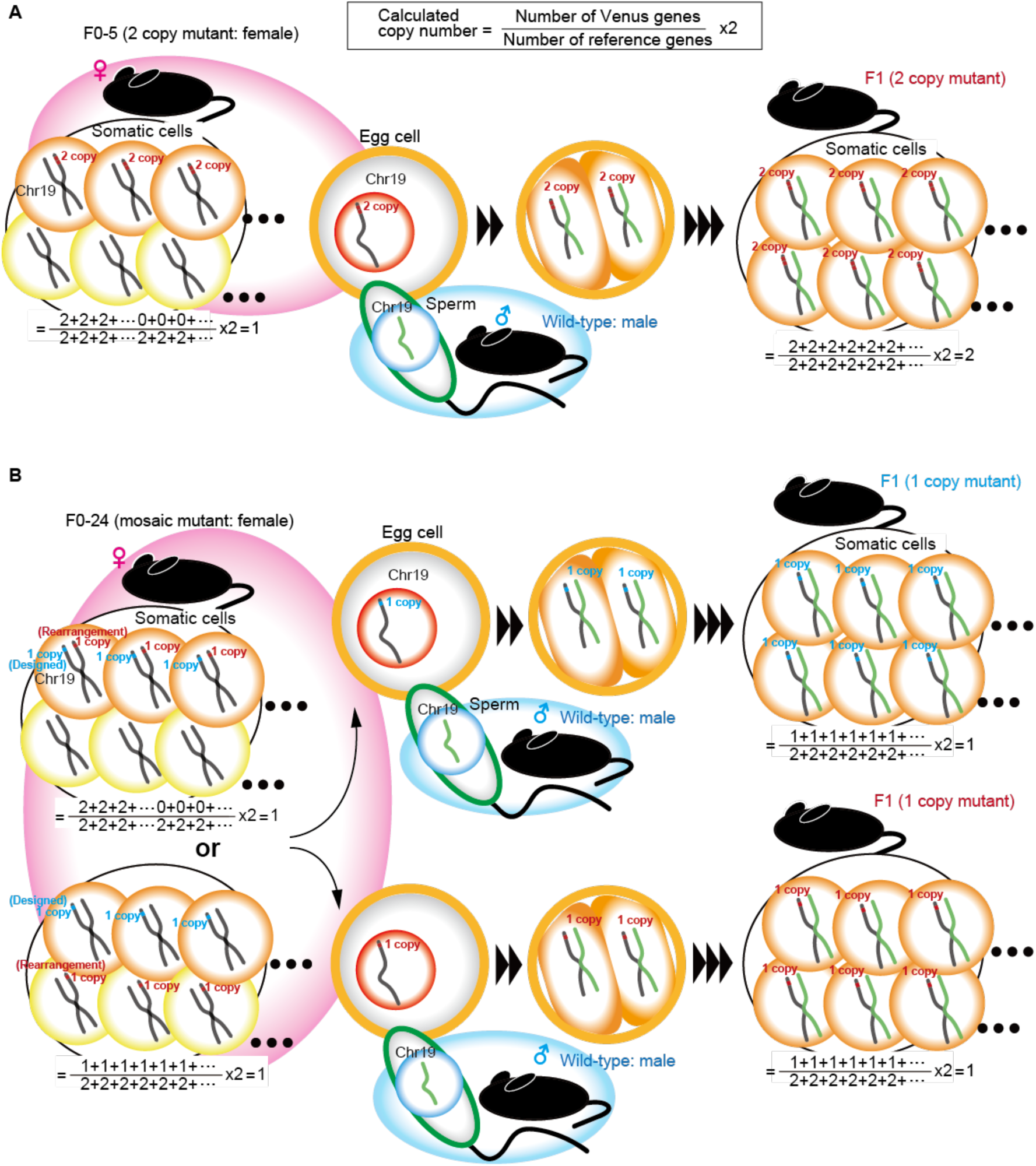
The designed knock-in sequence was compared to the PCR product 3’ ext-2 amplified from number 1 of the F1 generation. No unexpected mutations were detected in the two PCR products.

**Figure SI 9.** Schematic diagram comparing the calculated copy numbers of the F0 generation and the F1 generation obtained by crossing F0 with the wild type.

(A) Female F0 number 5 had an allele with a tandem two-copy insertion. The egg cell with the two-copy knock-in allele was fertilized by a wild-type sperm and the F1 generation was born. In F1 mice with two-copy alleles, somatic cells were homogeneous, and droplet digital PCR correctly detected them as two-copy. (B) Female F0 number 24 was a mosaic of a one-copy knock-in allele as designed and a one-copy knock-in allele in which genomic rearrengement occurred near the prlhr allele. Egg cells with each one copy knock-in allele were fertilized with wild-type sperm to produce the F1 generation. In F1 mice with one-copy alleles, somatic cells were homogeneous, and droplet digital PCR correctly detected them as one-copy.

Table S1. Results of Mapping with Magic-BLAST and annotation with SnpEff (Excel file)

**Table S2.**
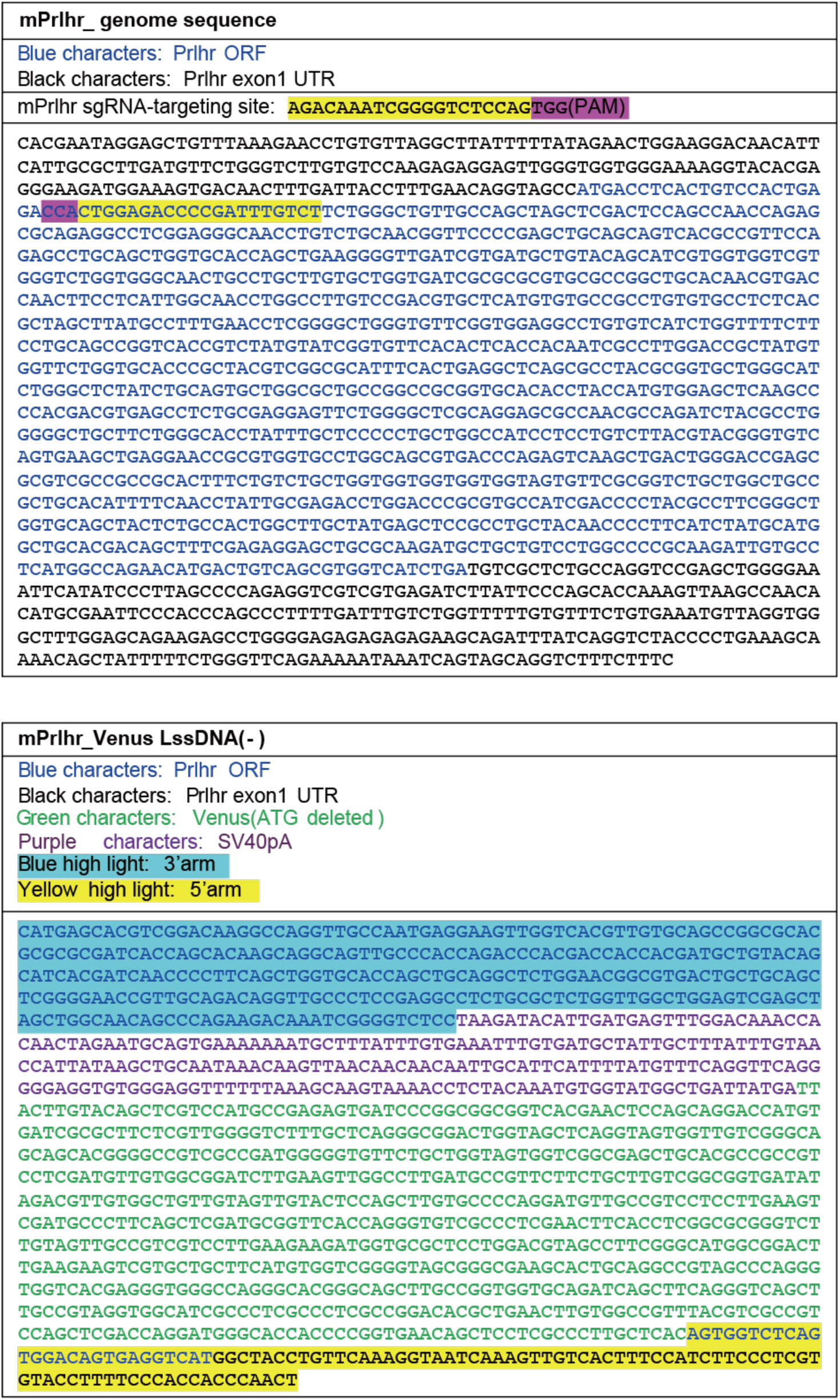
sgRNA designed to target the mouse Prlhr locus and sequence of donor DNA

Table S3. Primers (Excel file)

**Table S4.**
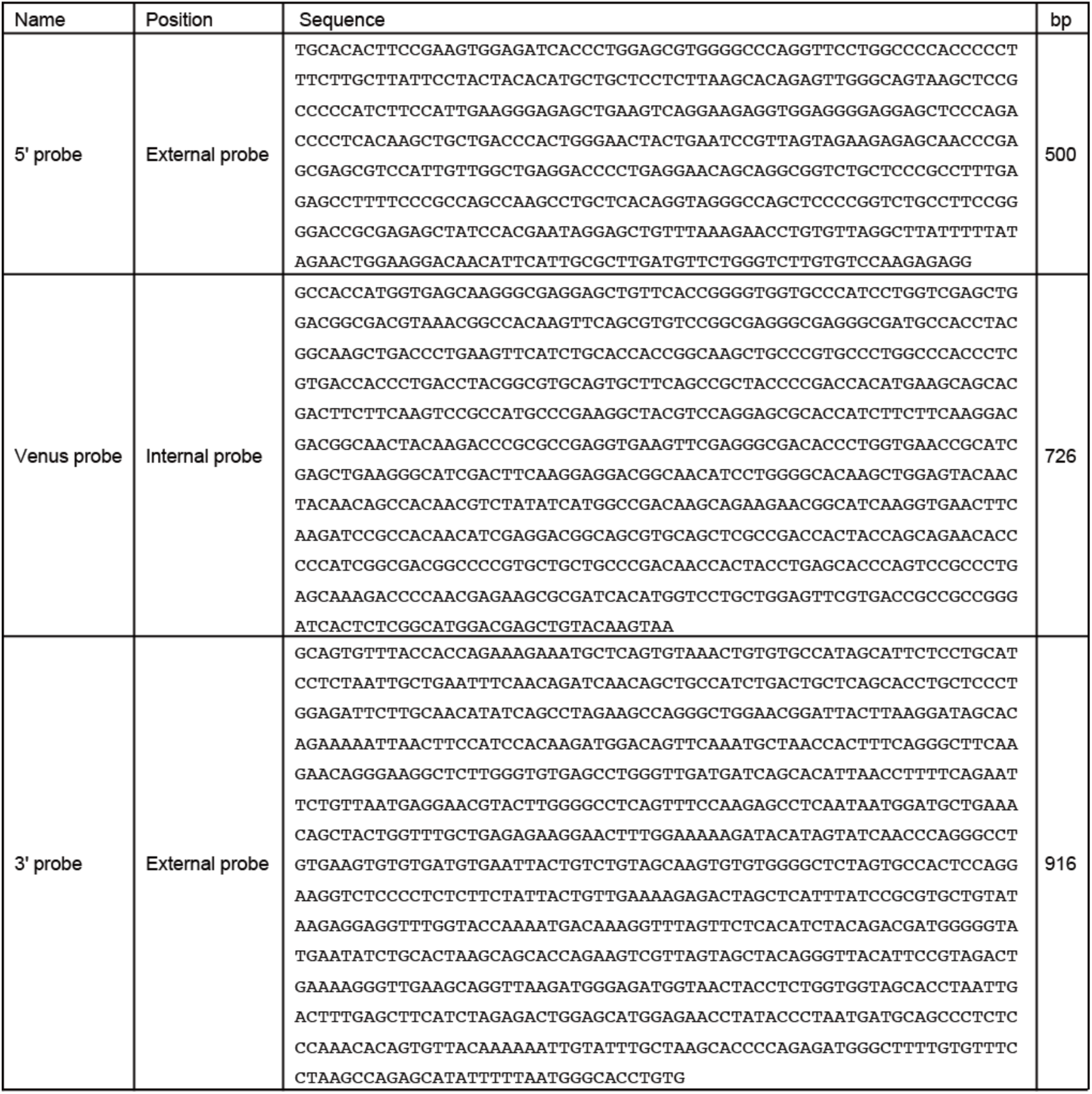
List of DNA probes used in Southern blot analysis

Table S5. Number of output reads from Miseq and number of reads after fastp processing (Excel file)

